# Constrained Transcriptional Polarity in the Organization of Mammalian *Hox* Gene Clusters

**DOI:** 10.1101/572875

**Authors:** Fabrice Darbellay, Célia Bochaton, Lucille Lopez-Delisle, Bénédicte Mascrez, Patrick Tschopp, Saskia Delpretti, Jozsef Zakany, Denis Duboule

## Abstract

In many animal species with a bilateral symmetry, *Hox* genes are clustered either at one or at several genomic loci. This organization has a functional relevance, as the transcriptional control applied to each gene depends upon its relative position within the gene cluster. It was previously noted that vertebrate *Hox* clusters display a much higher level of genomic organization than their invertebrate counterparts. The former are always more compact than the latter, they are generally devoid of repeats and of interspersed genes, and all genes are transcribed by the same DNA strand, suggesting that particular factors constrained these clusters towards a tighter structure during the evolution of the vertebrate lineage. Here we investigate the importance of uniform transcriptional orientation by engineering several alleles within the *HoxD* cluster such as to invert one or several transcription unit(s), with or without a neighboring CTCF site. We observe that the association between the tight structure of mammalian *Hox* clusters and their regulation makes inversions likely detrimental to the proper implementation of this complex genetic system. We propose that the consolidation of *Hox* clusters in vertebrates, including transcriptional polarity, evolved in conjunction with the emergence of global gene regulation *via* the flanking regulatory landscapes, to optimize a coordinated response of selected subsets of target genes in *cis*.

## INTRODUCTION

*Hox* genes are key players in the organization of the animal body plan. They encode transcription factors, the combination of which can instruct cells at different body levels as to their future morphological contributions. In addition to this ancestral function along the anterior to posterior axis, HOX proteins also participate in the organization either of secondary axes, or of a variety of organs or structures. In many animals displaying a bilateral symmetry, *Hox* genes are found clustered in the genome. This particular genomic topology has a functional relevance as the succession of genes in *cis* within the cluster corresponds to the order of their expression domains along the various axes. This collinearity phenomenon was initially proposed by Ed Lewis in *Drosophila* (Lewis 1978) and subsequently extended to vertebrates (Gaunt et al. 1988; Duboule and Dolle 1989; Graham et al. 1989; Akam 1989). In fact, *Drosophila* shows a breakpoint into the *Hox* cluster (Kaufman et al. 1980), which was thus split into two sub-clusters, a separation that occurred repetitively at different positions within drosophilids (Negre and Ruiz 2007). Also, some species display a complete disaggregation of their ancestral cluster into a collection of single gene loci, such as in Urochordata (Ikuta et al. 2004).

The existence of an entire *Hox* gene cluster, i.e. when all major paralogy groups are present and linked together in *cis*, was proposed to be always associated with animals whose segmental development occurs in a rostral to caudal time-sequence. In such animals, the activation of *Hox* genes must occur following a precise timing, referred to as the *Hox* clock, which would be either regulated or coordinated by the clustered organization (refs in (Duboule 2007). The availability of genome sequences for all major groups of animals has not yet proved this conjecture wrong. However, genome analyses have revealed an unexpected property for vertebrate *Hox* clusters, which differ from their invertebrate counterparts by a higher order in their structural organization. For instance, vertebrate *Hox* clusters are barely over 100Kb in size, whereas the cephalocordate or echinoderm clusters are ca. 500Kb large, similar to all characterized single invertebrate clusters (Fig. 1A). Therefore, the current situation is that not a single animal outside gnathostome (jawed) vertebrates has been reported to carry a complete *Hox* gene cluster of a size and compaction level close to that of vertebrates (Fig. 1B).

**Figure 1:**
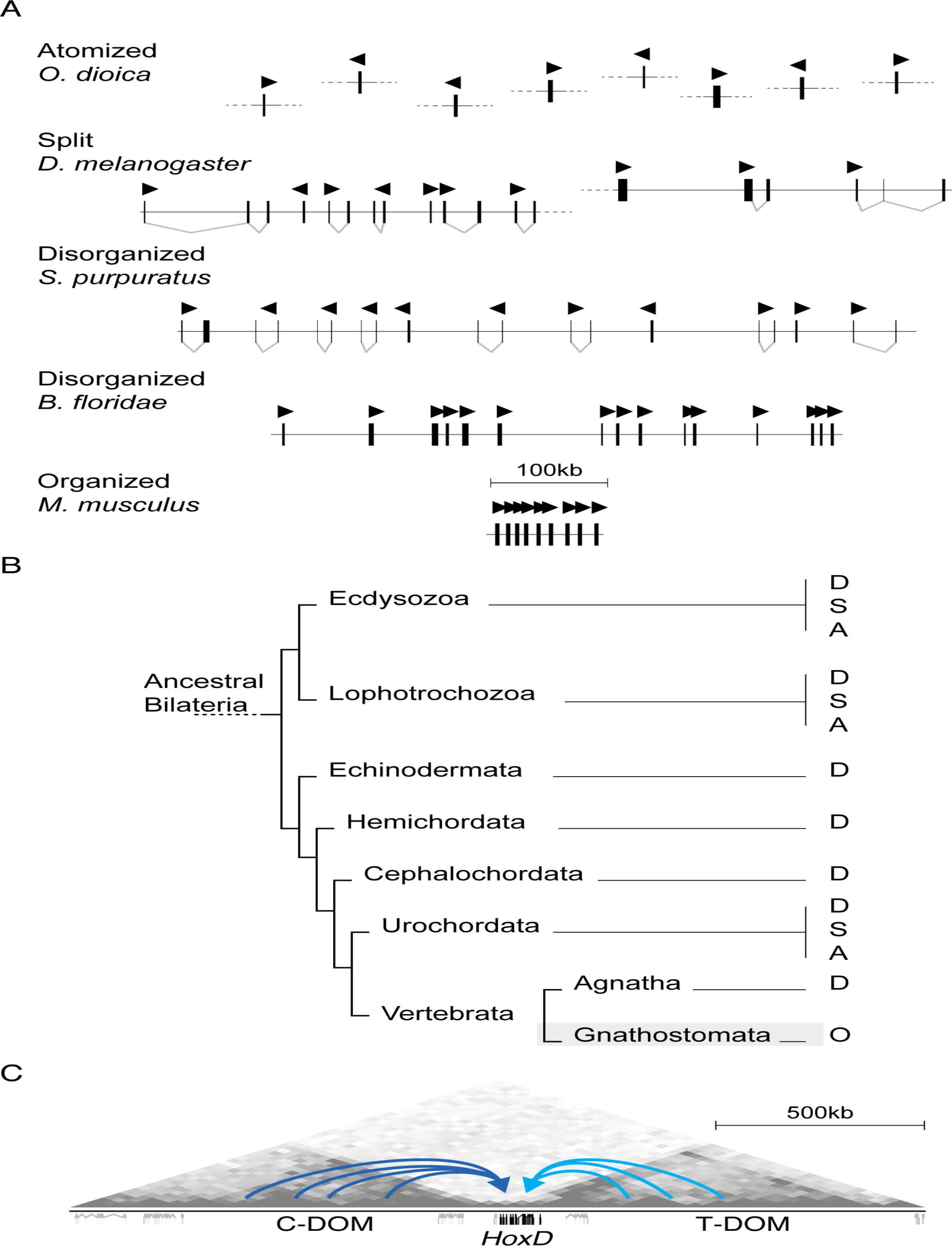
**A.** Structures of various *Hox* clusters drawn at the same scale. The highest level of organization and compaction is found in all jawed vertebrate genomes (type O, see (Duboule 2007). The arrowheads indicate the sense of *Hox* transcription. **B.** Atomized (A), split (S) or disorganized (D) *Hox* clusters can be found in other taxa (adapted from (Darbellay and Duboule 2016). **C.** Integration of the murine *HoxD* cluster at the border between two Topologically Associating Domains (TADs). Both of these TADs (C-DOM and T-DOM) contain a large gene desert including batteries of long-range enhancers (blue arrows) active at different developmental periods and in various structures (Hi-C data adapted from (Rodriguez-Carballo et al. 2017).

In addition to their general reduction in size, vertebrate *Hox* clusters contain little if any repeats, they never include genes unrelated to the *Hox* family and they transcribe all their genes from the same DNA strand, with a direction of transcription opposite to the time sequence of gene activation. Another important difference between the vertebrate genomes and those of cephalocordates, echinoderms and most invertebrates is that they contain multiple copies of their *Hox* clusters, as a result of the two initial rounds of genome duplication (2R, (Ohno 1970; Holland et al. 1994; Putnam et al. 2008). This exact correlation between the number of *Hox* gene clusters on the one hand, and their higher level of compaction and organization, on the other hand, lead to the proposal that these two distinct aspects may have co-evolved at the origin of the vertebrate lineage. A possible scenario was proposed whereby, much in the same way neo-functionalization can occur after horizontal or vertical gene duplication, global gene regulation achieved through the flanking regulatory landscapes may have been favored after duplication of the entire gene cluster (Figure 1C). In this view, the duplication of genomic *Hox* loci may have allowed the emergence of multiple potent enhancer sequences located outside the clusters and controlling several *Hox* genes at once. The potential negative effects of such high-order regulations and structures, for example upon the ancestral colinear mechanism at work during axial extension, could have been compensated for by having several clusters implementing this colinear process at the same time (Darbellay and Duboule 2016).

On the other hand, the evolution of *de novo* global enhancers may have represented an interesting adaptive value, in particular to evolve redundancy, compensatory mechanism or quantitative regulatory controls. As a result of this accumulation of enhancers in the regulatory landscapes flanking *Hox* clusters, the genes (or sub-groups thereof-) would have progressively maximized their responses to these enhancers, leading to a stepwise elimination of interfering repeat sequences, shortening of intergenic distances and placing all genes in the same transcriptional orientation. The latter point is of importance, for in such a tightly organized group of genes, the inversion of a transcription unit could either interfere with the neighboring genes’ transcription or bring two promoters close to one another leading to similar regulatory controls, a situation that would go against the general principle governing the evolution of this temporal mechanism in vertebrates. The presence of bound CTCF protein, a factor known to be involved in the insulation of chromatin domains (Ong and Corces 2014; Merkenschlager and Nora 2016) between almost every gene of the cluster (Soshnikova et al. 2010) supports the importance of this iterative genomic topology for a precise processing of gene activation.

In this study, we investigate the importance of the transcriptional directionality, in physiological conditions, by producing and analyzing a set of targeted inversions within the *HoxD* cluster and looking at the induced effects over the neighboring genes in various developmental contexts. We report the impact of inverting both the *Hoxd11* and *Hoxd12* loci, separately, without disturbing the distribution of intervening CTCF sites, as well as the effect of a combined inversion of the two loci together, along with the repositioning of an inverted CTCF site. While the former two inversions revealed regulatory disturbances, they lead to rather minor effects, whose long-term impact upon the animals were difficult to evaluate. The larger inversion, in contrast, elicited a dramatic up-regulation of the neighboring *Hoxd13* gene. By using additional engineered alleles, we show that this up-regulation is likely due to the reorganization of chromatin micro-domains, rather than the leakage of transcription on the opposite DNA strand, sent towards *Hoxd13* by the inverted *Hox* genes. We show that such chromatin domains are separated by a critical CTCF site, the mere deletion of which also leads to a transitory up-regulation of *Hoxd13* in the developing metanephric kidneys. Finally, we use a different allele to illustrate the deleterious effect of a stable gain-of-expression of *Hoxd13* in developing metanephros.

## RESULTS

### Inversion of the *Hoxd11* locus

By using the CRISPR/Cas9 technology, we produced a first inversion involving the *Hoxd11* transcription unit. The 5’ part of the *HoxD* cluster was selected due to the rather late timing of activation of these genes, which makes their study easier in terms of developmental stages, amount of material to collect and phenotypes to observe than their more 3’-located neighbors. This inversion was designed not to interfere with neighboring transcription start or termination sites and did not include any of the CTCF binding sites capable of recruiting this protein in those developmental contexts considered in this work (Fig. 2A). Therefore, the potential side-effects (i.e. any effect not generated by the mere inversion of *Hoxd11* transcription) were reduced to a minimum.

**Figure 2:**
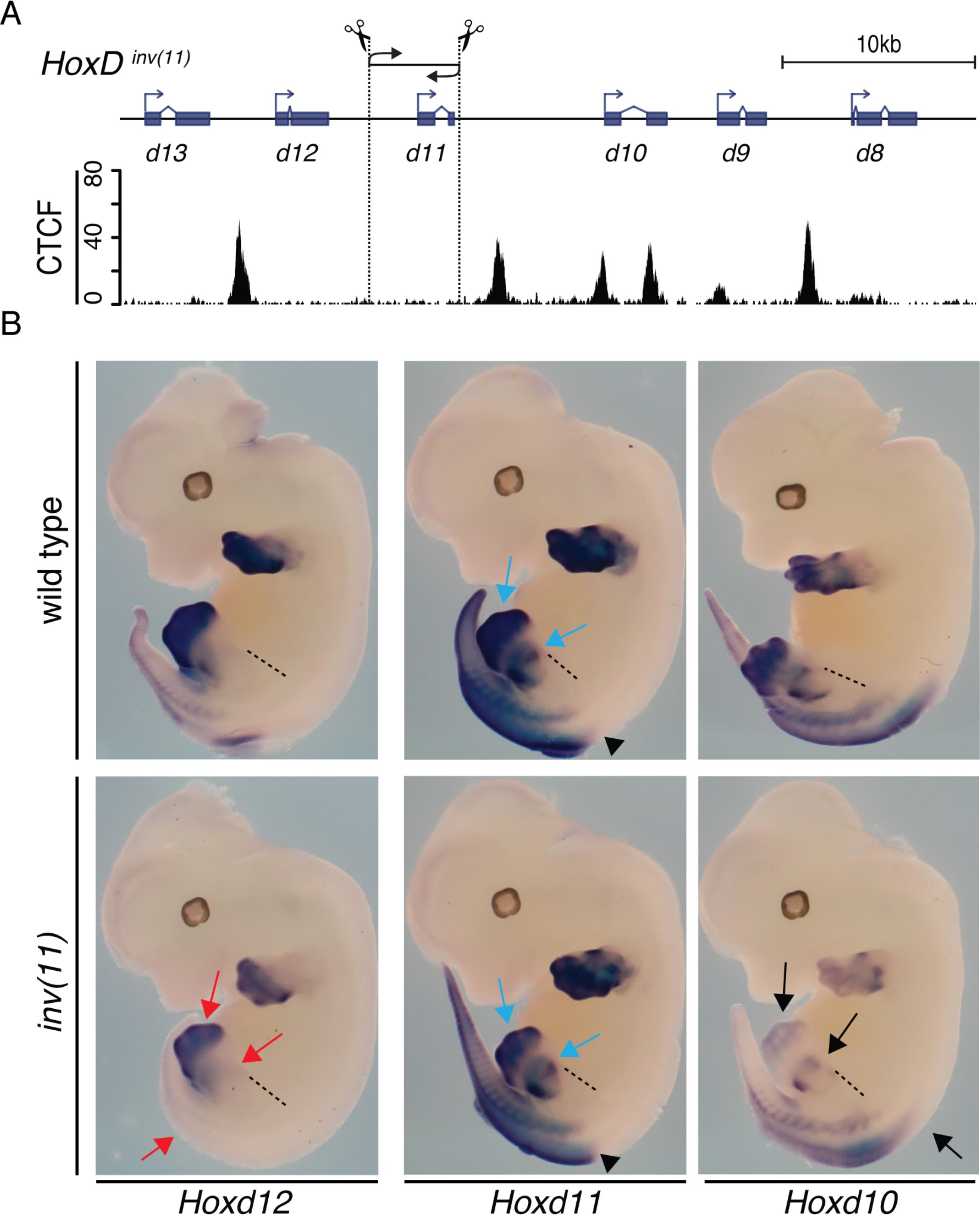
**A.** Scheme of the *HoxD*^*inv(11)*^ allele. The inversion is 4.6 kb in size. Below is a CTCF ChIPmentation profile from E13.5 wild-type metanephros. **B.** Whole mount *in-situ* hybridization of *Hoxd10*, *d11* and *d12* in E12.5 embryo. Homozygous *inv(11)* and wild-type littermates were used. The different expression patterns of *Hoxd11* are all preserved in the *inv(11)* allele (blue arrows and black arrowhead). In contrast the expression of both *Hoxd12* and *Hoxd10* is substantially reduced in the *inv(11)* mutant embryos (black and red arrows). In particular, *Hoxd12* is no longer detectable along the trunk axis, whereas the signal intensity is reduced both in the proximal and distal limb domains (red arrows). Dashed lines mark the anterior position of the hindlimb for reference.

Founder animals were produced and a strain referred to as *HoxD^inv(11)^* was derived and is mentioned as *inv(11)* mice or allele throughout this work. Animals carrying this inversion were first analyzed by whole mount *in-situ* hybridization (WISH) at embryonic day 12.5 (E12.5). In control wild-type animals of this stage, *Hoxd11* expression was well established both along the primary body axis, where it labeled the transition between the lumbar and the sacral regions (Fig. 2B, central panel on top, arrowheads), as well as the proximal and distal parts of developing limb buds (blue arrows). At the same stage, *Hoxd12* was expressed more posteriorly, whereas *Hoxd10* transcripts were found at more anterior positions, following the rule of collinearity in the expression of these genes (Fig. 2B, top left and right panels, respectively). This rule states that the more 5’-located is a gene positioned within its *Hox* cluster, the later this gene will be activated and the more posterior its expression domain will be.

WISH using homozygous *inv(11)* mutant littermates showed a clear decrease in the amount of detected *Hoxd10* and *Hoxd12* mRNAs visible in both the developing trunk axis and limb buds (Fig. 2B, bottom panels). Because WISH is not sufficiently quantitative, we carried out RNA-seq analyses by using two tissues, developing digits and metanephros, which were selected for several reasons. First, the two tissues strongly express *Hoxd11* under normal conditions and both require its function for proper development. Secondly, the two tissues express distinct combinations of other *Hoxd* genes, due to the topologies of their regulations. In presumptive digit cells, *Hoxd11* is transcribed along with *Hoxd10*, *Hoxd12* and *Hoxd13* as well as the *Evx2* gene positioned 8.8 kb upstream of *Hoxd13*. Altogether, these genes respond to series of enhancers located in the centromeric regulatory landscape (C-DOM), which coincides with a topologically associating domain (TAD)(Figure 1C)(Andrey et al. 2013). In contrast, future metanephric cells express *Hoxd11* together with *Hoxd10*, *Hoxd9, Hoxd8* and a moderate level of *Hoxd12* due to enhancers positioned in the opposite telomeric regulatory landscape (or T-DOM; Figure 1C), whereas *Hoxd13* mRNAs are not detected. While expression of *Hoxd13* and *Hoxd12* are necessary for digit development, they are detrimental to the development of metanephric kidneys due to a probable dominant-negative effect over other HOX proteins referred to as posterior prevalence (Duboule and Morata 1994).

In digit cells, *Hoxd13* is normally expressed the strongest, followed by *Hoxd12* and *Hoxd11* (Figure 3A, B). While the transcription termination seems to occur faithfully for *Hoxd13*, the RNA-seq dataset obtained from *HoxD*^*del(8-13)/+*^ control fetuses revealed transcriptional leakages of both *Hoxd12* and *Hoxd11*. These two genes exhibit low levels of mRNAs extending towards their neighboring downstream *Hox* transcription units, respectively *Hoxd11* and *Hoxd10* (Figure 3B, arrows). After inversion of *Hoxd11* (Figure 3C, shaded area), *Hoxd11* mRNAs, now encoded by the other DNA strand, continued at high level towards the *Hoxd12* transcription unit (Figure 3C, middle anti-*Hox* profile, arrow). This transcription leakage was likely due to the fact that the major termination signal for *Hoxd11* was not inverted along with the transcription unit. This abnormally elongated *Hoxd11* transcript extended up to the termination site of the *Hoxd12* mRNA, where it decreased abruptly, as shown by the superimposition of both DNA strands (Figure 3C, bottom profile, arrowhead). However, some *Hoxd11* transcripts continued to leak over the *Hoxd12* locus up to the 3’ extremity of *Hoxd13* (Figure 3C, bottom profile, grey arrowhead). In parallel, transcription of *Hoxd12*, from the other DNA strand, was greatly reduced (Figure 3A, arrow), as if the abnormal level and extension of *Hoxd11* transcription on the anti-*Hox* coding strand would interfere with the correct transcription process of *Hoxd12* from the normal -non-inverted-strand (Figure 3C, top and bottom profiles).

**Figure 3:**
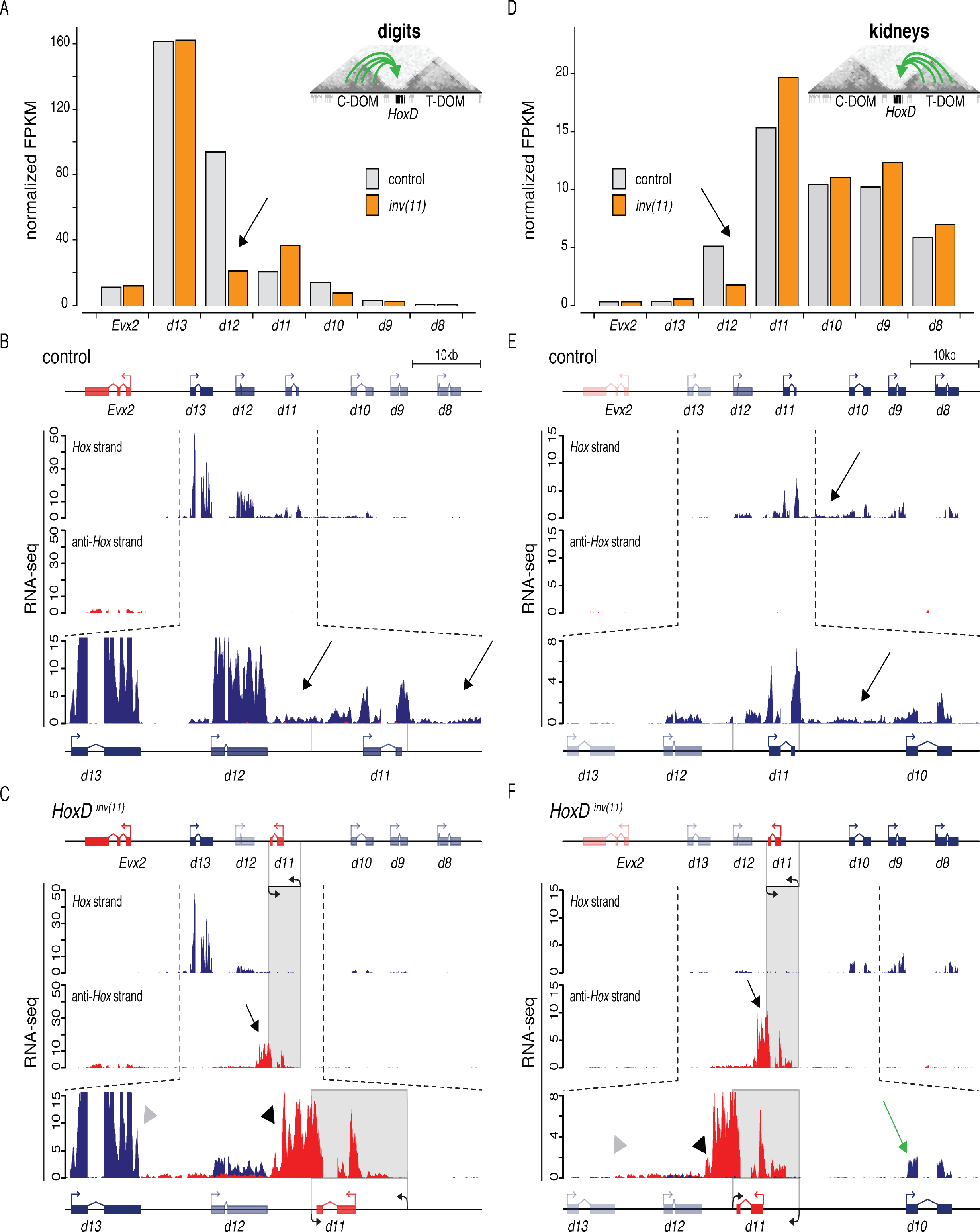
**A.** Normalized quantification (FPKM) of *Hoxd* RNAs in E12.5 digits in control *HoxD*^*del(8-13)/+*^ (grey) and *HoxD*^*del(8-13)/inv(11)*^ mutant (orange). *Hoxd* genes display quantitative collinearity with *Hoxd13* expressed the strongest (Montavon, 2008). In the *inv(11)* allele, *Hoxd12* RNAs are reduced (black arrow). **B.** Strand specific RNA signals from heterozygous control *HoxD*^*del(8-13)/+*^ developing digits. Both *Hoxd12* and *Hoxd11* transcripts leak towards their 3’ neighbor gene (black arrows). **C.** Strand specific RNA signals from trans-heterozygous *HoxD*^*del(8-13)/inv(11)*^ developing digits. Inversion of *Hoxd11* (shaded area) resulted in transcripts leakage towards *Hoxd12* (arrows), which stopped downstream of *Hoxd12* (black arrowhead). Some transcripts extended towards *Hoxd13*, whose transcription remained unaffected until its 3’ end (grey arrowhead). **D.** Normalized quantification (FPKM) of *Hoxd* RNAs in E13.5 metanephros. *Hoxd12* expression in the *inv(11)* allele also decreases (black arrow), despite its low level in control metanephros due to a posteriorly restricted domain. **E.** Strand specific RNA signals from control heterozygous *HoxD*^*del(8-13)/+*^ E13.5 metanephros. *Hoxd8* to *Hoxd11* are highly transcribed whereas *Hoxd13* and *Evx2* are silent. Some *Hoxd11* transcripts leak over *Hoxd10* (black arrow). **F.** Strand specific RNA signals from trans-heterozygous *HoxD*^*del(8-13)/inv(11)*^ mutant metanephros. Leakage of *Hoxd11* transcripts on the anti-*Hox* DNA strand was maintained (black arrow). A fraction of these transcripts extended up to *Hoxd13* 3’ UTR termination site (grey arrowhead) and thus covered *Hoxd12*. *Hoxd13* remained silent in this mutant allele. The inversion of *Hoxd11* does not reduce the amount of *Hoxd10* transcripts (green arrow). The signals in B, C, E and F were normalized by the number of million uniquely mapped reads. The mapped RNA signals are shown either in blue (*Hox* DNA strand) or in red (anti-*Hox* DNA strand). The inverted *Hoxd11* locus is represented by a shaded area. The vertical grey lines around *Hoxd11* in the control scheme (last track of B and E) indicate the breakpoints of the inverted allele.

In contrast to digit cells, developing metanephric cells do not express any *Hoxd13* mRNAs and only moderate levels of *Hoxd12*, with *Hoxd11*, *Hoxd10* and *Hoxd9* being the most expressed genes (Figure 3D). As in digit cells, however, the inversion of the *Hoxd11* locus led to a decrease in the level of *Hoxd12* mRNAs (Figure 3D). Once again, the *Hoxd11* transcripts now encoded by the *Hox* opposite DNA strand did not terminate after exon 2, as in the normal case (Figure 3E, arrowhead), but extended up to the termination site of the *Hoxd12* transcripts (Figure 3F, black arrow and arrowhead), with some weak but clear signal continuing over the *Hoxd12* transcription unit. Of note, in the inverted configuration, *Hoxd10* transcripts were scored as in the wild-type chromosome, showing that under normal conditions, the transcription of this gene is not dependent upon the leakage of transcripts coming from *Hoxd11* and extending up to *Hoxd10* (Figure 3E, black arrow).

Therefore, in this particular case, the inversion of a *Hox* gene locus induced the down-regulation of transcripts coming from the gene located immediately in 5’, likely as a result of a collision effect of the transcriptional machineries associated with the respective genes. It is unclear as to why the opposite effect was not scored, i.e. why the transcription of the inverted *Hoxd11* gene was not affected by *Hoxd12* transcripts, in particular in digit cells where *Hoxd12* is normally expressed at higher levels than *Hoxd11* and where the former gene seems to also send some transcripts towards the latter (Figure 3B, arrows). It is nevertheless clear that *Hoxd11* transcripts have the capacity to negatively interfere with *Hoxd12* transcription, whereas the opposite was not observed.

### Inversion of the *Hoxd12* locus

We next inverted the *Hoxd12* transcription unit, i.e. the piece of DNA just adjacent to the *Hoxd11* 5’ inversion break point, but still leaving in place the CTCF site separating these two loci from the *Hoxd13* gene (Figure 4A). As for the inversion of *Hoxd11*, we looked at the effect of this inversion on the transcription profiles of both developing metanephros and digits. RNA quantification of a subset of *Hoxd* genes in E13.5 metanephros revealed opposite variations in the amount of *Hoxd12* and *Hoxd11* mRNA, with an increase of *Hoxd12* transcription while *Hoxd11* was slightly decreased (Figure 4B, black arrows). The expression of other *Hoxd* genes remained unchanged. This increase was equally visible on the RNA-seq profiles obtained from *HoxD*^*del(8-13)/inv(12)*^ trans-heterozygote mutant metanephros (Figure 4C, D, compare red and blue profiles). The inversion of *Hoxd12* also resulted in a limited transcriptional leakage from the anti-*Hox* DNA strand, extending towards *Hoxd13* (Figure 4C, black arrow). However, transcripts terminated at the same position as previously observed for the *inv(11)* allele (Figure 3C, F, Figure 4C, grey arrowhead) and hence they did not overlap with the *Hoxd13* gene. *Hoxd13* and *Evx2* remain transcriptionally silent.

**Figure 4:**
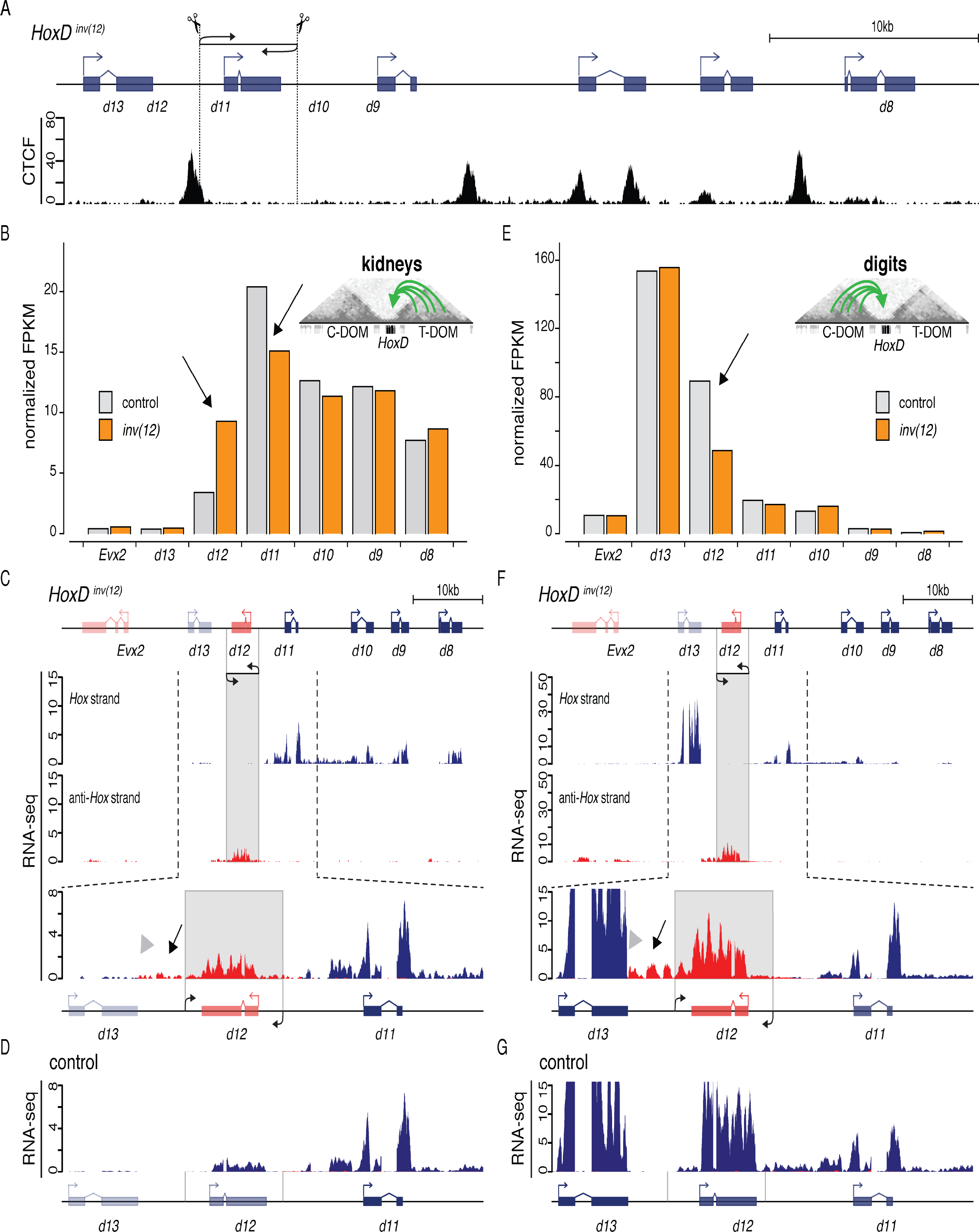
**A.** Scheme of the CRISPR/Cas9 guides used to engineer the *HoxD*^*inv(12)*^ allele. Below is a CTCF ChIPmentation profile from E13.5 wild-type metanephros. **B.** Normalized quantification (FPKM) of *Hoxd* genes RNAs in E13.5 metanephros in control *HoxD*^*del(8-13)/+*^ (grey) and mutant *HoxD*^*del(8-13)/inv(12)*^ (orange). In the *inv(12)* allele, the amounts of both *Hoxd12* and *Hoxd11* mRNAs are modified (black arrows). **C.** Strand specific RNA signals from trans-heterozygous *HoxD*^*del(8-13)/inv(12)*^ mutant metanephros. A weak leakage of *Hoxd12* transcripts was scored (black arrow), which stopped at *Hoxd13* 3’ UTR termination (grey arrowhead, see also Fig. 3C, F). *Hoxd13* and *Evx2* remained silent. **D.** Strand specific RNAs obtained from *HoxD*^*del(8-13)/+*^ E13.5 metanephros (adapted from Figure 3B). **E.** Normalized quantification (FPKM) of *Hoxd* genes RNAs in E13.5 developing digits. In this allele, the amount of *Hoxd12* RNAs is reduced (black arrow), other *Hoxd* mRNAs remained constant. **F.** Strand specific RNA signals from trans-heterozygous *HoxD*^*del(8-13)/inv*(*12*)^ mutant developing digits. The leaking *Hoxd12* RNAs stop 3’ of *Hoxd13*, at the same position as in panel C (black arrow). **G.** Strand specific RNA signals from control heterozygous *HoxD*^*del(8-13)/+*^ digits (adapted from Figure 3E). *Hoxd13* transcription efficiently terminates and does not leak towards *Hoxd12*. The signals used in C, D, F and G were normalized by the number of million uniquely mapped reads. The mapped RNA signals are shown either in blue (*Hox* DNA strand) or in red (anti-*Hox* DNA strand). The inverted *Hoxd12* locus is represented by a shaded area. The vertical grey lines around *Hoxd12* in the control schemes (D and G) indicate the breakpoints of the inverted allele.

During digit development, the amount of *Hoxd12* transcripts was reduced in *inv(12)*, in contrast to the situation described in metanephros, whereas other *Hoxd* genes did not display any substantial variation in RNA content (Figure 4E). The RNA-seq profiles obtained from developing trans-heterozygous *HoxD^del(8-13)/inv(12^*^)^ mutant digits confirmed the minimal impact, if any, of this inversion on the transcription of *Hoxd13* (Figure 4F, G). The transcriptional leakage coming from the inverted *Hoxd12* locus abruptly terminated 3’ of *Hoxd13* (Figure 4F, black arrow and grey arrowhead), at the same position already reported for the *inv(12)* allele in metanephros (Figure 4C). Interestingly, in both cases, this transcript went through the position of the bound CTCF and terminates at the polyA site of *Hoxd13*, even when this latter gene was transcribed barely over background (Figure 4C, F).

### Inversion of both *Hoxd11* and *Hoxd12*

The independent inversions of either the *Hoxd11*, or the *Hoxd12* loci did not significantly disturb the general transcriptional activity within the *HoxD* cluster. Importantly, both inverted DNA segments were contained between two bound CTCF sites, without perturbing either their locations or their orientations. Therefore, we looked in detail at an inversion of both *Hoxd11* and *Hoxd12* in *cis*, whereby the CTCF site located between *Hoxd12* and *Hoxd13* (Figure 5A, black arrow) was inverted along with the two transcription units (Kmita et al. 2000). In this *HoxD*^*inv(11-12)d13lacZ*^ allele, a *lacZ* cassette was introduced in-frame to the first exon of *Hoxd13*, which was thus functionally inactivated.

**Figure 5:**
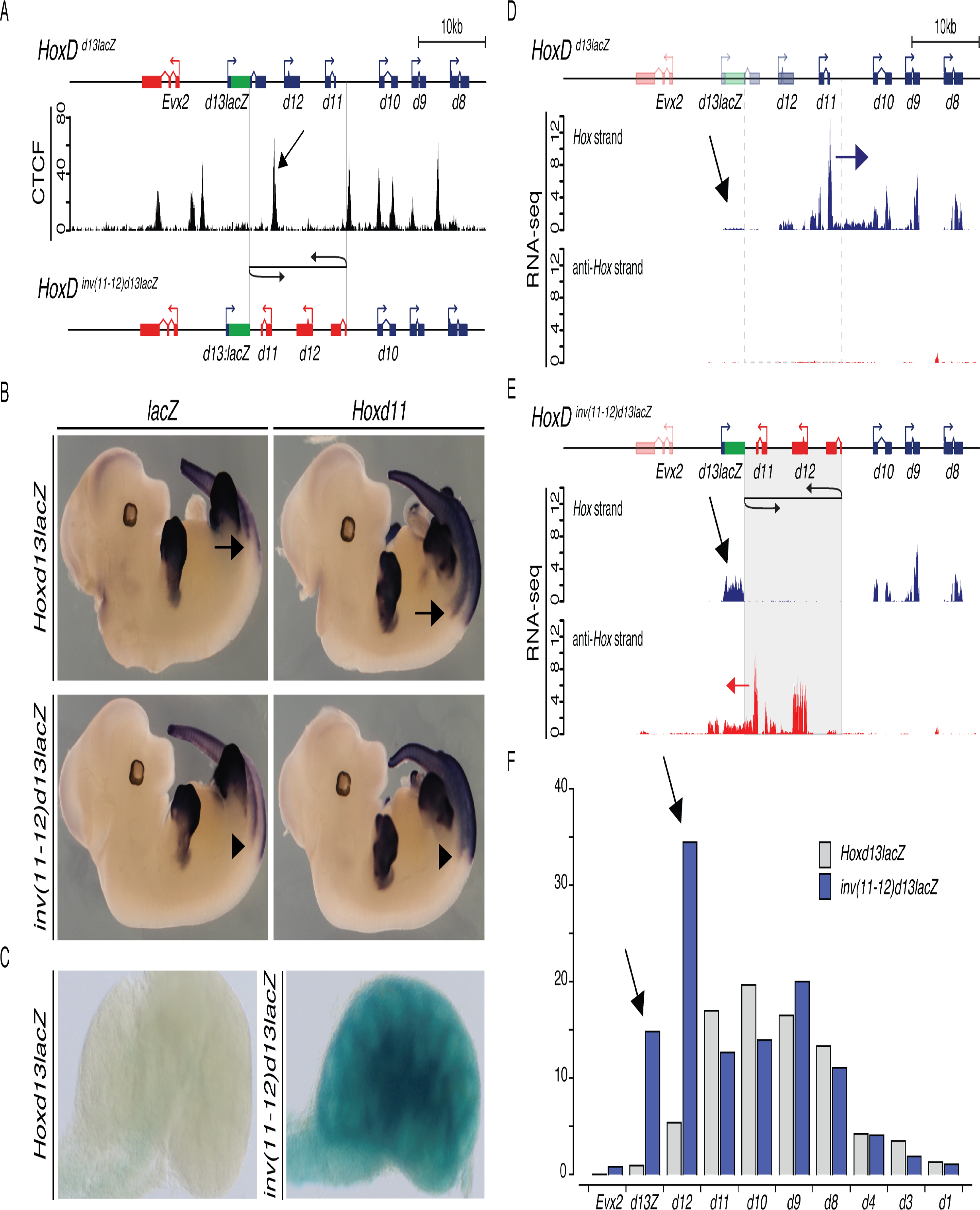
**A.** Schemes of the *HoxD*^*d13lacZ*^ and *HoxD*^*inv(11-12)d13lacZ*^ alleles (see also (Kmita et al. 2000). In between is a CTCF ChIPmentation profile from E13.5 metanephros of *HoxD*^*d13lacZ*^ embryos. In the *HoxD*^*inv(11-12)d13lacZ*^ allele, both *Hoxd11* and *Hoxd12* were inverted as well as the CTCF site positioned between *Hoxd12* and *Hoxd13lacZ* (black arrow). **B.** WISH of *lacZ* and *Hoxd11* in control *HoxD*^*d13lacZ*^ and *HoxD*^*inv(11-12)d13lacZ*^ E12.5 homozygous mutant embryos. In the control, the rostral limits of *Hoxd11* and *d13lacZ* are well separated along the AP axis (black arrows). In the *inv(11-12)d13lacZ allele*, these patterns merged due to the anteriorized expression of *Hoxd13lacZ* (black arrowheads). **C.** X-gal staining of E13.5 metanephros from homozygote *HoxD*^*d13lacZ*^ (left) and *HoxD*^*inv(11-12)d13lacZ*^ embryos (right). Staining is scored in *HoxD*^*inv(11-12)d13lacZ*^ metanephros only. **D.** Strand specific RNA profiles from homozygous *HoxD*^*d13lacZ*^ E13.5 metanephros. The reporter *Hoxd13lacZ* was not transcribed (black arrow), whereas *Hoxd12* showed a low RNA amount. RNAs leaking from *Hoxd11* to *Hoxd10* are shown (blue arrow). The dotted square represents the region which is inverted in the *HoxD*^*inv(11-12)d13lacZ*^ allele. **E.** Strand specific RNAs from homozygous *HoxD*^*inv(11-12)d13lacZ*^ E13.5 metanephros. *Hoxd11* transcripts now leak over *Hoxd13lacZ* (red arrow) concomitantly with the ectopic transcription of *Hoxd13lacZ* (see panel B). **F.** Normalized quantification (FPKM) of *Hoxd* RNAs before and after inversion. *Hoxd13lacZ* and *Hoxd12* RNAs were gained (black arrows). Signals in D and E were normalized by the number of million uniquely mapped reads. The mapped RNA signals are shown either in blue (*Hox* DNA strand) or in red (anti-*Hox* DNA strand). The inverted *Hoxd11-d12* locus is highlighted by a shaded area. The *lacZ* sequence knocked-in *Hoxd13* is shown as a green box.

We compared the WISH signal before (*HoxD*^*d13lacZ*^) and after (*HoxD*^inv(*11-12*)*d13lacZ*^) inversion and noticed an anteriorization of the staining indicating a gain-of-expression of *Hoxd13lacZ* in more anterior trunk territories in the inverted allele (Figure 5B, left panels, arrow and arrowhead). In the inverted mutant allele, *Hoxd13lacZ* was expressed at an anterior level now equivalent to that of *Hoxd11* (Figure 5B. bottom panels, arrowhead), which did not substantially change after inversion (Figure 5B, right panels, arrow and arrowhead). *HoxD*^*d13lacZ*^ fetuses did not show any X-gal staining in developing metanephros (Figure 5C, left), in agreement with the absence of *Hoxd13* transcription normally observed in this organ located rostral to *Hoxd13* expression boundary. In contrast, a robust staining was scored in *HoxD*^inv(*11-12*)*d13lacZ*^ metanephros, in conjunction with the general anteriorization of *Hoxd13* expression (Figure 5C, right).

RNA-seq profiles obtained from homozygous *HoxD*^*d13lacZ*^ E13.5 metanephros confirmed the quasi-absence of *Hoxd13lacZ* transcripts (Figure 5D, black arrow), with the majority of mRNAs produced by the central *Hoxd11* to *Hoxd8* genes, all from the same *Hox* DNA strand, whereas *Hoxd12* showed a low level of activity. An important leakage of *Hoxd11* RNAs extending towards *Hoxd10* was detected (Figure 5D, blue arrow). After inversion, several changes were observed. First, the transcription of *Hoxd12*, now at the relative genomic position of former *Hoxd11*, was increased. Secondly, *Hoxd11* transcription was maintained at a rather high level, despite its new relative position matching that of former *Hoxd12*, and its transcripts now leaked towards the *Hoxd13lacZ* gene, though on the opposite anti-*Hox* DNA strand (Figure 5E, red arrow). Concomitantly, *Hoxd13lacZ* was significantly up-regulated in metanephros from inverted mutant fetuses, even though its relative position in the gene cluster had not changed (Figure 5E, black arrow). These differences in RNA amounts were quantified by computing the FPKM values, confirming the gain of signal for both the *Hoxd13lacZ* and *Hoxd12* genes, located on opposite DNA strands (Figure 5F, black arrows).

This ectopic activation of *Hoxd1*3 was accompanied by modifications in both the coverage in some histone marks and the micro-3D structure of the locus. In the control locus, PRC2-dependent H3K27me3 marks (Figure 6A, orange profile), which label silent genes, were particularly enriched over both *Hoxd13* and *Evx2*, the two genes completely inactive during metanephros development (Figure 6A, arrowed bracket). Other *Hoxd* genes were also labeled by this mark, though to a lesser extent, due to the presence of mixed cellular populations in this developing organ including a fair proportion of *Hoxd*-negative cells. The transition between high and low levels of H3K27me3 marks coincided with the presence of both a bound CTCF site and a RAD21 peak (Figure 6A, large blue arrowhead). This CTCF site labels the TAD boundary at the *HoxD* cluster (Figure 6A, vertical dashed line and (Rodriguez-Carballo et al. 2017)) and is positioned exactly where clusters of CTCF sites change their orientations (Figure 6A, blue and red arrowhead). Therefore, it seemed that domains of high *versus* low H3K27me3 coverage were separated by the first CTCF site with an orientation (blue) opposed to all CTCF sites but one found in the center of the cluster (red). This transition in histone marks was reinforced by the analysis of H3K4me3 modifications (Figure 6A, green profile), which displayed a distribution complementary to those of H3K27me3 marks with a robust coverage over *Hoxd9* to *Hoxd11*, a weaker signal over *Hoxd12* and only traces over *Hoxd13lacZ*.

In the inverted allele, the H3K27me3 and H3K4me3 profiles remained complementary (Figure 6B, track 2, orange and green respectively). Yet, the boundary was clearly displaced towards the centromeric end of the gene cluster. The amount of H3K27me3 over *Hoxd13lacZ* was severely reduced when compared to the coverage over *Evx2* (Figure 6B, bracketed arrowheads) while at the same time H3K4me3 marks robustly increased over the *Hoxd13lacZ* gene. The boundary between these two complementary epigenetic profiles was now positioned between *Hoxd13lacZ* and *Evx2* (Figure 6B, dashed line) and precisely matched the presence of a pair of CTCF sites orientated in the direction of the centromeric TAD (circled in Figure 6B). The CTCF site that labeled this transition in the control allele (Figure 6A, large blue arrowhead) no longer marked the transition in the inverted allele (Figure 6B, large red arrowhead), even though this site was still occupied by CTCF and matched a peak of RAD21 in the mutant allele (Figure 6B, arrow in track 3).

This shift in the position of the boundary between active and inactive domains of the *HoxD* cluster in the inverted allele was challenged by a 4C-seq approach whereby the contacts established by the *Hoxd13lacZ* gene (Figure 6A, B, track 4) were assessed before and after the inversion. In the control allele, contacts were enriched towards the centromeric side, i.e. in agreement with the position of the boundary (Figure 6A, track 4). In the inverted allele, however, the bulk of contacts were now detected towards the telomeric side, i.e. following the change in the position of the boundary (Figure 6B, track 4). Therefore, the inversion importantly modified the tropism in contacts of the *Hoxd13lacZ* gene, from contacts mainly established with a repressive chromatin structure before the inversion, to a clear enrichment of contacts with a transcriptionally permissive chromatin domain after the inversion. In the latter case, interestingly, the gain of contact appeared somehow restricted to the DNA interval between the new boundary (Figure 6B, dashed line) and the position of the inverted CTCF site (Figure 6B, large red arrowhead), as if a boundary effect was also observed at this position (Figure 6B, open arrowhead).

This effect was confirmed by using a viewpoint corresponding to *Hoxd9*, on the same material. In the control *HoxD*^*d13lacZ*^ allele, *Hoxd9* established contacts with all genes up to the CTCF-labeled boundary, where interactions abruptly stopped (Fig. 6A, bottom track). In the mutant *HoxD*^*inv(11-12)d13lacZ*^ allele, however, contacts extended up to the new boundary (Fig. 6B, bottom track) corresponding to the CTCF/RAD21 doublet located between *Hoxd13lacZ* and *Evx2* (circled in Fig. 6B). The new boundary effect triggered by the inverted CTCF site and observed with the *Hoxd13lacZ* viewpoint (Fig. 6B, open arrowhead) was nevertheless also detected when using *Hoxd9* as bait. Therefore, after inversion, while *Hoxd9* interacted with *Hoxd13lacZ*, these contacts appeared to result from the interactions between two micro sub-domains rather than from a single domain (Fig. 6B, bottom track).

**Figure 6:**
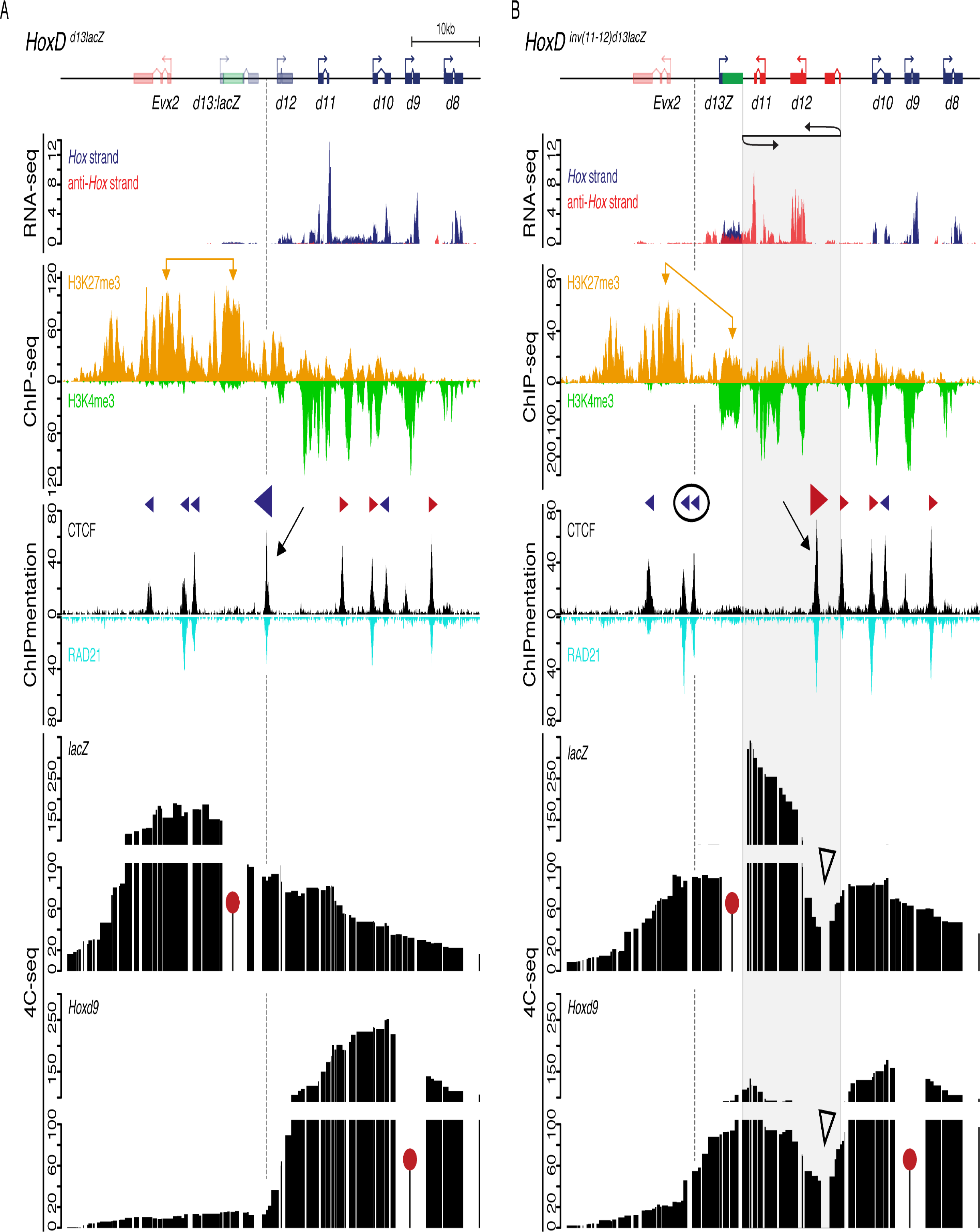
Chromatin marks, CTCF/RAD21 and 4C-seq profiles generated from either control homozygous *HoxD*^*d13lacZ*^ (**A**) or homozygous *HoxD*^*inv(11-12)d13lacZ*^ mutant (**B**) E13.5 metanephros. From top to bottom: The schemes of the alleles, strand specific RNA (adapted from Figure 5D and 5E), H3K27me3 (orange) and H3K4me3 (green) profiles, CTCF (black) and RAD21 (turquoise blue) profiles and 4C-seq profiles using either a *lacZ* (top) or a *Hoxd9* (bottom) viewpoint (red pins). The H3K4me3 and H3K27me3 profiles reveal two domains within the *HoxD* cluster with silent genes in 5’ and active genes in 3’. **A.** In control metanephros, both *Hoxd13* and *Evx2* show high levels of H3K27me3 marks (bracketed orange arrows) and little if any H3K4me3 marks, with a boundary matching a CTCF/RAD21 peak (black arrow and vertical dashed dark line), which display an orientation facing the C-DOM (large blue arrowhead). 4C-seq profiles reveal that *Hoxd13lacZ* mostly contacted the 5’ domain, while *Hoxd9* had an opposite interaction tropism, respecting the position of the CTCF site (vertical dashed line). **B.** Comparable tracks from homozygous *HoxD*^*inv(11-12)d13lacZ*^ E13.5 mutant metanephros. From top to bottom: normalized strand specific mRNA RNA-seq (adapted from Figure 5E), ChIP-seq of H3K27me3. The inversion leads to a gain of *Hoxd13lacZ* expression along with a diminution of H3K27me3 over *Hoxd13* (bracketed yellow arrows) and a gain of H3K4me3 thus repositioning the boundary between *Evx2* and *Hoxd13* (vertical dashed line). This position coincided with a pair of CTCF sites (circled), which then labelled the new centromeric side of the boundary, while the initial CTCF site (large arrowhead in A) was inverted and displaced (large red arrowhead). The latter site still binds CTCF and RAD21 (black arrow). The 4C-seq profiles after inversion were in agreement with the position of this new boundary. The *lacZ* bait had more interaction on the telomeric than on the centromeric side and the *Hoxd9* bait now extended its contacts up to include *Hoxd13lacZ* to stop at the new boundary. The inverted CTCF site induced a slight and local boundary effect (open arrowheads). The inverted DNA segment is shown with a shaded area. Green box: *lacZ* sequences.

### Change in chromatin topology or promoter cleaning?

These results suggested that local changes in chromatin topology, due to the modified position and orientation of a CTCF site, were responsible for the up-regulation of the *Hoxd13lacZ* gene in developing metanephros. Alternatively, the transcript emanating from inverted *Hoxd11* and overlapping with the *Hoxd13lacZ* gene on the opposite anti-*Hox* DNA strand could elicit a transcriptional response from the latter unit. Such ‘promoter cleaning’ would allow subsequent control by the appropriate upstream factors, which would normally not access this promoter due to the coverage by H3K27me3 marks. To try to discriminate between these two explanations, we further deleted the *Hoxd11* transcription units from the inverted allele such as to abrogate the RNA leakage over *Hoxd13lacZ*. In this *HoxD*^*inv(11-12)del(11)d13lacZ*^ allele, only *Hoxd12* was left inverted and positioned at the same distance from *Hoxd13lacZ* than inverted *Hoxd11* in the *HoxD*^*inv(11-12)d13lacZ*^ allele. This secondary mutation did not affect the distribution of CTCF sites in any way (Figure 7A, top).

**Figure 7:**
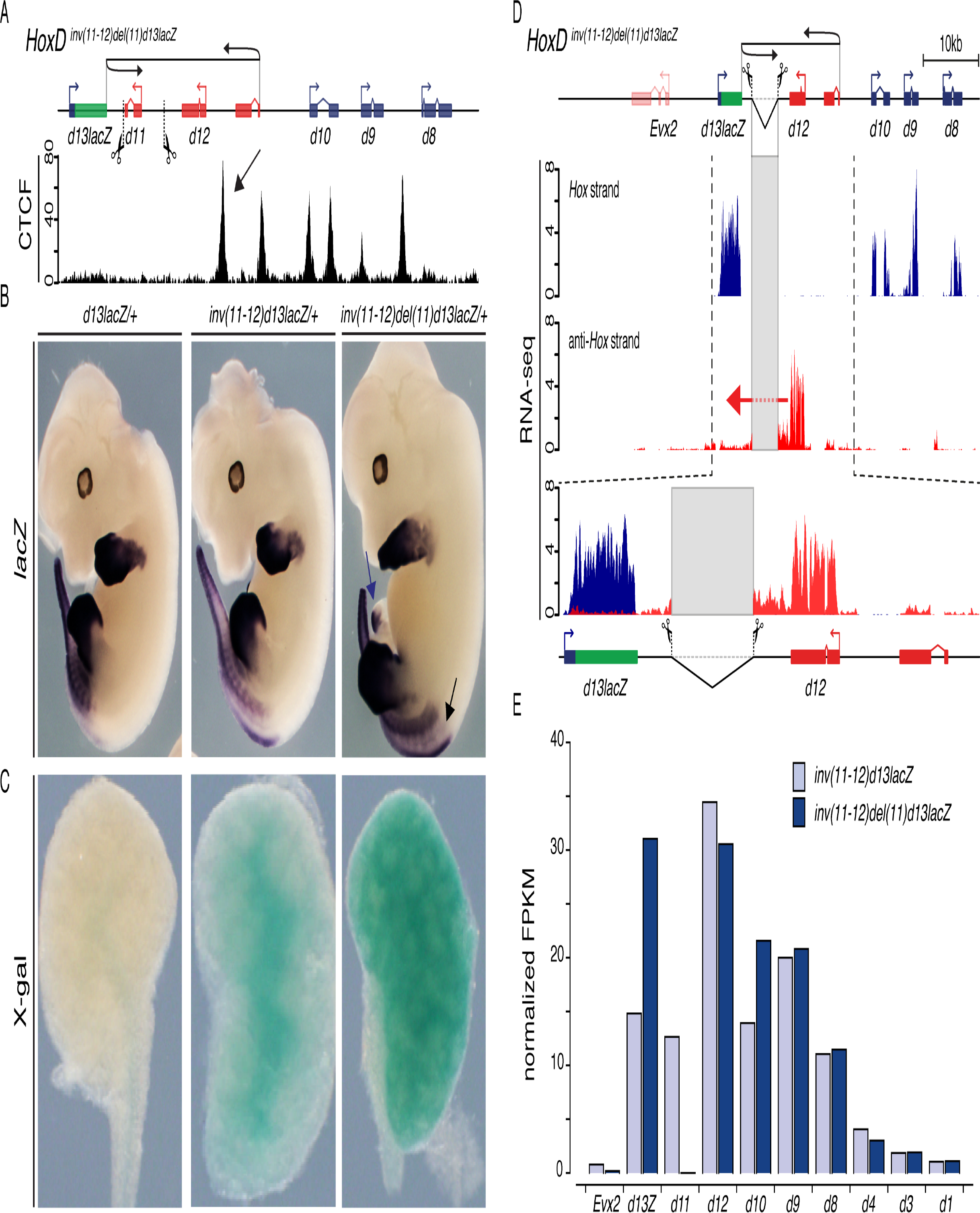
**A.** Scheme of the CRISPR/Cas9 guides used to engineer the *HoxD*^*inv(11-12)del(11)d13lacZ*^ allele. Below is a CTCF ChIPmentation profile from metanephros of E13.5 *HoxD*^*inv(11-12)d13lacZ*^ homozygous embryos (adapted from Figure 6B). The *CTCFd12d13* site is inverted and repositioned as in Figure 6B (black arrow). **B.** WISH using a *lacZ* probe of *d13lacZ* (left), *inv(11-12)d13lacZ* (middle) and *inv(11-12)del(11)d13lacZ* (right) E12.5 heterozygous embryos. A gain of *d13lacZ* expression in the caecum and along the main body axis is observed when compared to the *d13lacZ* control (black arrowhead and arrow, respectively). **C.** X-gal staining of E13.5 metanephros carrying the same alleles as in panel B. Staining is scored in the heterozygous *inv(11-12)del(11)d13lacZ* metanephros indicating that *d13lacZ* remains transcriptionally active in this mutant. **D.** Strand specific RNAs from homozygous *HoxD*^*inv(11-12)del(11)d13lacZ*^ E13.5 metanephros. Few *Hoxd12* transcripts (red) extend towards the *d13lacZ* gene (red arrow). **E.** Normalized quantification (FPKM) of *Hoxd* transcripts. The important gain of *d13lacZ* in the *inv(11-12)del(11)d13lacZ* allele, where *Hoxd12* transcript leakage towards the former locus is minimal suggests that the former event does not depend on the latter. The signals in D were normalized by the number of million uniquely mapped reads. The deleted *Hoxd11* locus is highlighted by a shaded area. *lacZ* sequences are shown by a green box.

WISH using a *lacZ* probe revealed that the deletion of *Hoxd11* from the *inv(11-12)d13lacZ* allele failed to suppress the anteriorization of *Hoxd13lacZ* along the AP axis of *HoxD*^*inv(11-12)d13lacZ*^ fetuses (Figure 7B, black arrow). This indicated that the anteriorization was likely not due to *Hoxd11* transcripts cleaning the *Hoxd13lacZ* promoter and induced a *lacZ* pattern similar to that of inverted *Hoxd11* (Figure 7B, middle and right panels). Also, *HoxD*^*inv(11-12)del(11)d13lacZ*^ mutant fetuses transcribed *lacZ* in their developing caecum (Figure 7B, blue arrow), similar to their aged-matched *HoxD*^*inv(11-12)d13lacZ*^ counterparts (see also (Kmita et al. 2000; Delpretti et al. 2013), suggesting again that this ectopic expression was not caused by a *Hoxd11* transcript leakage. Likewise, X-gal staining was scored in E13.5 *HoxD*^*inv(11-12)del(11)d13lacZ*^ metanephros, similar to- or even stronger than the *HoxD*^*inv(11-12)d13lacZ*^ situation (Figure 7C).

Strand specific RNA-seq carried out using E13.5 homozygous *HoxD*^*inv(11-12)del(11)d13lacZ*^ metanephros revealed the expected transcriptional activity of *Hoxd12* on the opposite strand as well as the strong gain-of-expression of the *Hoxd13lacZ* reporter gene (Figure 7D). A faint *Hoxd12* transcript was detected extending up to the *Hoxd13* promoter, yet this RNA was in much lower amount than the leaking *Hoxd11* RNA observed in the *HoxD*^*inv(11-12)d13lacZ*^ allele (compare Figures 7D and 5E, red arrows). We compared this decrease in amount of leaking RNAs observed between the *HoxD*^*inv(11-12)del(11)d13lacZ*^ and the *HoxD*^*inv(11-12)d13lacZ*^ alleles, with a quantification of the genes’ transcripts obtained by computing the FPKM described above. The steady-state levels of the various mRNA were globally comparable between the two alleles, yet with a clear increase in *Hoxd13lacZ* mRNAs in the *HoxD*^*inv(11-12)del(11)d13lacZ*^ allele (which expectedly also lacked any *Hoxd11* transcripts)(Figure 7E). This latter observation suggested a negative correlation between the amount of *Hoxd13lacZ* mRNA produced, on the one hand, and the importance of transcript leaking on the opposite DNA strand, on the other hand.

### Mutation of the CTCF site

As a consequence of these results, a change in local topology was favored over a mere transcriptional leakage, as a cause for the up-regulation of the *Hoxd13lacZ* reporter gene. Since the CTCF site present in the inverted DNA seemed to play a particular function in the spatial organization of these chromatin domains, we set out to mutate this site in a control genetic background. This *HoxD*^*del(CTCF:d12d13)*^ allele was generated by CRISPR/Cas9-induced mutagenesis using a gRNA targeting the core of the CTCF binding motif (Fig. 8, pink sequence), positioned between *Hoxd12* and *Hoxd13.* The selected deletion was 21 bp in size, including 13 bp of the core CTCF motif. CTCF ChIPmentation in trans-heterozygote *HoxD*^*del(CTCF:d12d13*)/*del*(*8-13)*^ E13.5 forebrain confirmed the total absence of CTCF binding at this mutated site (Fig. 8B, black arrows), whereas other CTCF peaks remained unchanged.

**Figure 8:**
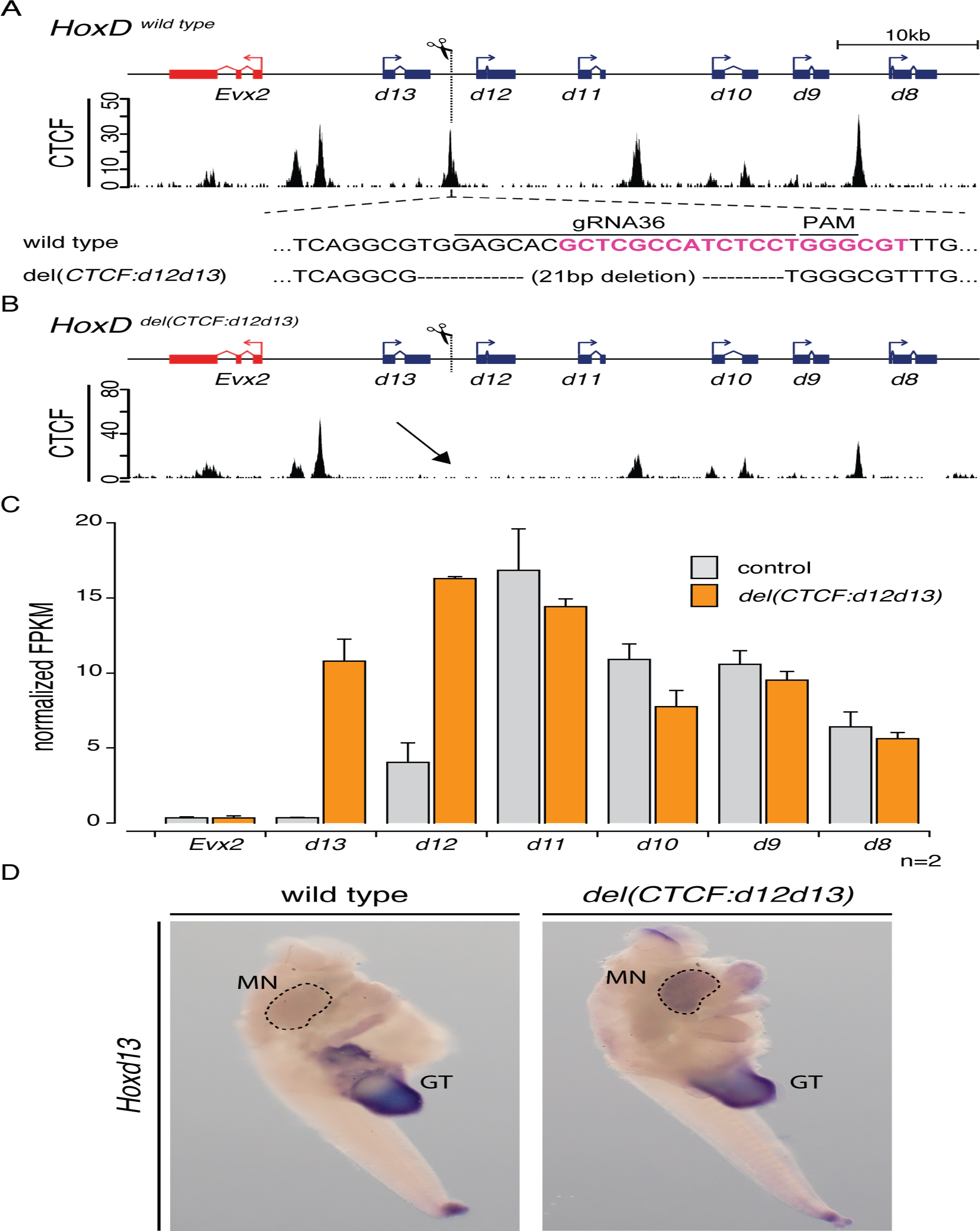
**A.** Scheme of the *HoxD*^*del(CTCF:d12d13*)^ allele. The core of the CTCF binding motif targeted by gRNA36 (20bp) is shown in pink. The del(*CTCF:d12d13*) allele corresponds to a deletion of 21 bp, 13 of which are within the CTCF motif. This CTCF profile is from E13.5 wild-type forebrain. **B.** The ChIPmentation of CTCF in *HoxD*^*del(CTCF:d12d13)/del(8-13)*^ E13.5 forebrain cells shows that the binding of CTCF is abolished (black arrow), whereas other peaks remained globally unchanged. **C.** Normalized quantification (FPKM) of *Hoxd* RNAs from either *HoxD*^*del(CTCF:d12d13)/del(8-13)*^ or control *HoxD*^*del(8-13)/+*^ metanephros at E13.5. The gain-of-expression of *Hoxd12* and *Hoxd13* in *del(CTCF:d12d13)* metanephros is robust, whereas the neighboring *Evx2* gene remained transcriptionally silent. **D.** *In situ* hybridization of *Hoxd13* in E13.5 embryos. The abdominal cavity was opened and metanephros exposed before processing. Dissected posterior parts are shown. The gain-of-expression of *Hoxd13* in developing metanephros (MN) of the homozygous *del(CTCF:d12d13)* mutant is visible whereas *Hoxd13* is not expressed in a control wild-type littermate. *Hoxd13* signals are scored in the genital tubercle (GT) and the tip of the elongating tail bud in both mutant and wild-type embryos.

Dissected metanephros of E13.5 *HoxD*^*del(CTCF_d12d13)/del(8-13)*^ and control *HoxD*^*+/del(8-13)*^ fetuses were analyzed for their content in *Hoxd* mRNAs through normalized quantification (FPKM) computed from the RNA-seq datasets. In the absence of the CTCF site, a conspicuous gain-of-expression was scored for both *Hoxd12* and *Hoxd13*, i.e. those two *Hoxd* genes normally located on the other side of the mutated CTCF site, as if the boundary effect had entirely disappeared (Fig. 8C). This effect was further controlled by *in situ* hybridization using *Hoxd13* as a probe in E13.5 embryos, which revealed a clear gain-of-expression of *Hoxd13* in the developing metanephros (MN) of homozygous *HoxD*^*del(CTCF:d12d13)*^ mutant embryos (Fig. 8D).

This gain-of-function effect observed after the mutation of a single CTCF site was rapidly compensated for, however, and ectopic *Hoxd13* was already much reduced at E18.5 and no longer scored in adult metanephros of mutant specimens, whereas *Hoxd8* and *Hoxd9* transcripts were still scored at high levels in both control and mutant adult metanephros (Fig. S1). This result indicated that while the CTCF mutation had fragilized the boundary, the remaining CTCF sites as well as the transcription of other *Hoxd* genes might still exert a strong insulation effect, preventing HOXD13 protein to leak into the developing metanephros. Therefore, to address this question, we used another allele, where the gain of *Hoxd13* in metanephros was slightly more robust and, most importantly, more stable over time such that it was still scored in adult metanephros.

### Detrimental effect of HOXD13 on metanephric kidney development

To address this point, we used the *HoxD*^*inv(Nsi-Itga6)*^ allele, where an inversion was engineered directly upstream of *Hoxd13* up to the *Itga6* gene, three megabases away (Tschopp and Duboule 2011)(Fig. 9A, top). As a consequence of this inversion, the *Hoxd13* locus loses some of its strong contact points in the centromeric neighborhood and reallocates contacts towards its telomeric side (Andrey et al. 2013), where metanephros enhancers are located (Di-Poi et al. 2007). We looked by RNA-seq at whether this partial change in contact tropism may elicit a gain-of-function during metanephros development and a substantial gain was observed for *Hoxd13* transcripts while, at the same time, the levels of *Hoxd9*, *Hoxd10* and *Hoxd11* transcripts seemed to decrease, likely due to competition between promoters (Fig. 9B).

**Figure 9:**
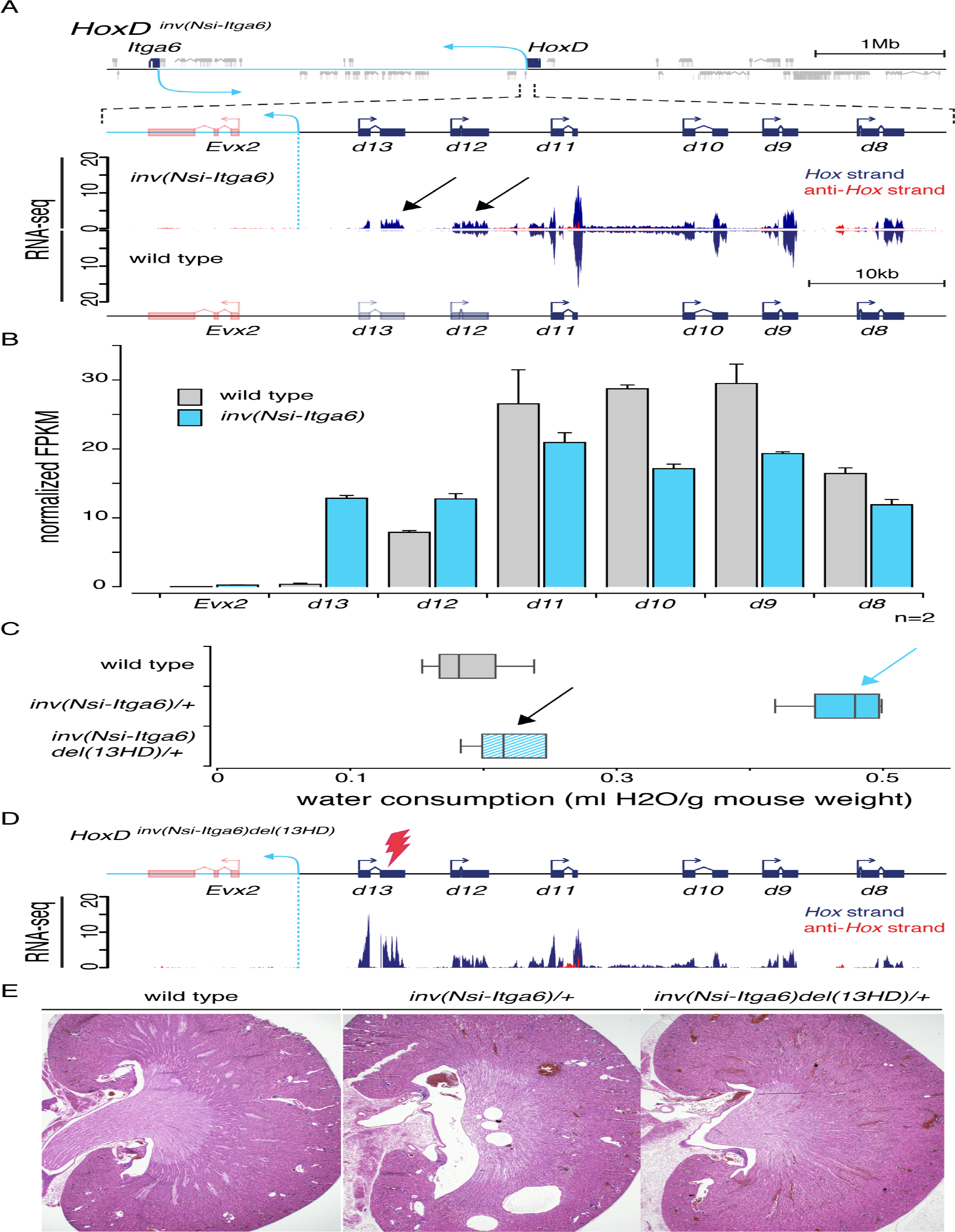
**A.** Gain of *Hoxd13* expression in the metanephros of *HoxD*^*inv(Nsi-Itga6)*^ mutant specimens. The *HoxD*^*inv(Nsi-Itga6)*^ allele, also referred to as *inv(Nsi-Itga6)*, contains a 3 Mb large inversion centromeric from the *HoxD* cluster (Tschopp and Duboule 2011) with a breakpoint positioned between *Evx2* and *Hoxd13* (top, dashed blue line). In E13.5 metanephros, a noticeable gain-of-expression of both *Hoxd12 and Hoxd13* genes is scored (black arrows). The mutant heterozygote *inv(Nsi-Itag6)* (top) and wild-type control (bottom) RNA profiles are shown in mirror orientations. **B.** Normalized quantification (FPKM) of *Hoxd* mRNAs computed from RNA-seq datasets obtained from heterozygous *HoxD*^*inv(Nsi-Itga6)/+*^ and wild-type E13.5 metanephros. **C.** Normalized quantification of the daily water intake in adult animals of different genotypes: wild type (top), *HoxD*^*inv(Nsi-Itga6)/+*^ (middle) and *HoxD*^*inv(Nsi-Itga6)del(13HD)/+*^(bottom). *HoxD*^*inv(Nsi-Itga6)*^ heterozygotes show a two-fold increased consumption of water (blue arrow) as compared to wild-type animals. The deletion of the *Hoxd13* homeodomain in this latter allele (*HoxD*^*inv(Nsi-Itga6)del(13HD)*^, see Supplementary Figure 2) rescues this phenotype (black arrow). **D.** RNA-seq profile from E13.5 metanephros micro-dissected from homozygous *HoxD*^*inv(Nsi-Itga6)del(13HD)*^ embryos. **E.** Representative sections of metanephros dissected from adult animal of different genotypes: wild type (left), *HoxD*^*inv(Nsi-Itga6)*^ heterozygote (middle) and double mutant *HoxD*^*inv(Nsi-Itga6)del(13HD)*^ heterozygote. Alterations are only detected in *HoxD*^*inv(Nsi-Itga6)*^ mutant specimen.

Mice carrying one copy of this allele were found to suffer from polydipsia, i.e. they were consuming more than twice the amount of water than their control littermates (Fig. 9C, blue arrow). A histological analysis of heterozygous mutant metanephros revealed serious malformations, in particular at the level of the medulla, suggesting that the polydipsia was indeed due to problems in metanephric kidney development (Figure 9E, middle panel). The (*Nsi-Itga6*) inversion that caused this phenotype is large and has a breakpoint in a gene rich region around the *Itga6* locus and hence the contribution of the gained HOXD13 protein in this kidney alteration remained to be clearly established.

To this aim, we used a CRISPR/Cas9 approach to induce a second mutation editing further the inversion allele, which would inactivate the function of the HOXD13 protein (Fig. 9D, top). We induced a small deletion into the homeobox region of HOXD13 (the *HoxD*^*inv(Nsi-Itga6)del(13HD)*^ allele, see Supplementary Figure 2) and verified that the transcript profile displays similar features, in particular with respect to the gain of *Hoxd13* transcripts in developing metanephros (Fig. 9D). Mice carrying this double-mutant rescue allele displayed a normal level of water consumption (Fig. 9C, black arrow) as well as metanephros with a normal morphology (Fig. 9E, right panel), indicating that inactivation of the homeodomain of ectopically expressed HOXD13 protein was able to revert the gain-of-function phenotype induced by the inversion. This experiment formally demonstrated that an anterior expansion of *Hoxd13* transcripts, in physiological amounts, was detrimental to the development of structures located anterior to its normal expression level, thus providing a selective pressure for the evolution and maintenance of a strong boundary effect in the *HoxD* cluster.

## DISCUSSION

Vertebrate *Hox* gene clusters display both a gene density and a level of organization thus far unmatched across metazoans, suggesting that a consolidation process occurred along with the evolution of the vertebrate lineage. This counter-intuitive conclusion (Duboule 2007) was tentatively explained by the emergence and implementation of global and long-range regulatory controls (Darbellay and Duboule 2016), which was itself made possible by the two rounds of genome amplification that occurred at the root of vertebrates (Ohno 1970; Holland et al. 1994). In this view, *Hox* genes became progressively more tightly organized to better respond to these remote regulations, a process that allowed the emergence of a coordinated regulation and lead to functional complexity and redundancy. This transition from a group of genes located in *cis* to a ‘meta-gene’ is materialized by a series of structural hallmarks such as the absence of foreign genes interspersed, the unusual scarcity of repeated elements in these gene clusters and their reduced sizes when compared to invertebrate *Hox* clusters. In this study, we challenged the functional relevance of yet another hallmark of these loci, i.e. the fact that all HOX proteins are encoded by the same DNA strand.

### Inversions with various effects

The various inversion alleles analyzed had different impacts upon the regulation of their neighbor *Hox* genes. While the inversion of *Hoxd12* had little (if any) effects upon either *Hoxd13* or *Hoxd11* regulation, the inversion of *Hoxd11* had a robust negative outcome upon the transcription of *Hoxd12*. In contrast, the inversion of both *Hoxd11* and *Hoxd12* in *cis* elicited a substantial gain-of-expression of *Hoxd13* in various developing tissues. While the functional impact of a down-regulation of *Hoxd12* has not yet been uncovered (Kondo et al. 1996; Davis and Capecchi 1996), a gain-of-expression of *Hoxd13* in places where it is normally not expressed leads to severe conditions, due to the dominant negative effect of the HOXD13 protein over other HOX products. In limbs and metanephros for instance, ectopic expression of *Hoxd13* was shown to correlate with severe anomalies (Herault et al. 1997; van der Hoeven et al. 1996), which in metanephros phenocopied the combined loss of function of *Hoxa11* and *Hoxd11* (Wellik et al. 2002). Therefore, the *Hoxd11-Hoxd12* inversion would very likely be detrimental to the mouse. In the current study, however, the inactivation of *Hoxd13* by a *lacZ* insertion prevented these phenotypes to develop and thus allowed for a detailed analysis to be carried out.

The transcriptional effects observed in these inversion alleles can be interpreted within two distinct, yet not exclusive, explanatory frameworks. The first one involves problems directly associated with the transcription dynamics of these genes, i.e. the necessity due to the high concentration of transcription units, to have all elongations occurring with the same polarity. In the wild-type situation, some transcription units do not stop before reaching the 5’ UTR of the gene located in 3’. For example, some *Hoxd11* transcripts extend over *Hoxd10* and, likewise, *Hoxd4* and *Hoxd3* exons are intermingled thus giving the possibilities for hybrid transcripts to be generated. Upon inversion, these transcription units may either collide with transcripts coming from the other DNA strand or trigger transcription of a silent gene through a ‘promoter cleaning’ effect.

The second explanatory framework involves modifications in the topology of the locus, which may directly impact upon the regulations of the various genes. For example, an inversion may bring a promoter close to that of the 3’ located neighbor gene, thus inducing promoter sharing and hence the mis-regulation of at least one of the two units. Alternatively, the inversion can reposition structural elements such as CpG islands or CTCF sites, which might in turn modify the general architecture of the locus, inducing novel enhancer-promoter contacts.

### Transcriptional interference of topological modification?

The decrease in the steady state level of *Hoxd12* transcripts upon *Hoxd11* inversion may reflect a collision effect, with transcripts elongating from *Hoxd11* and reaching the *Hoxd12* 3’ UTR, thus interfering with the correct transcription of this latter gene and its mRNA’s stability. In *S. cerevisiae*, the main source of transcriptional interference is caused by the frontal collision between two transcriptional machineries moving on opposed DNA strand (Hao et al. 2017; Pannunzio and Lieber 2016; Palmer et al. 2011). These collisions result in mutual arrest of the two complexes, leading to their dissociation from the DNA through polyubiquitylation of the RNA Pol II and the degradation of the aborted RNA transcripts (Hobson et al. 2012; Osato et al. 2007; Prescott and Proudfoot 2002; Somesh et al. 2005). It is however not known whether factors such as the relative elongation rates or some characteristics of the chromatin can selectively impact one RNA (Boldogköi 2012; Hao et al. 2017). In the case of the *Hoxd11-Hoxd12* inversion, however, transcripts elongating from *Hoxd11* did not collide with those of *Hoxd13lacZ* since the latter gene was silent and covered by H3K27me3 marks. This suggested that such *Hoxd11* transcripts were able to displace these repressive chromatin marks from the *Hoxd13* promoter, thus allowing it to be transcribed in the same domains where *Hoxd11* was active.

Recent studies have highlighted the capacity of different PRC2 components to bind nascent RNA transcripts, resulting in some case in the eviction of PRC2 out of the chromatin and to the modulation of its EZH2-dependent methyl-transferase activity (Cifuentes-Rojas et al. 2014; Davidovich et al. 2013; Herzog et al. 2014; Kaneko et al. 2014). To demonstrate this possibility, we deleted the *Hoxd11* gene from this inverted allele such as to stop the mRNA leakage over *Hoxd13*. This was successfully achieved as the now repositioned inverted *Hoxd12* gene only sent few transcripts over the *Hoxd13lacZ* locus, in a much lower amount than inverted *Hoxd11* before its deletion. Yet the up-regulation of *Hoxd13lacZ* in metanephros was at least as strong as in the presence of inverted *Hoxd11*, if not stronger, suggesting that such a promoter cleaning mechanism was likely not to operate.

### Modifications in local chromatin topology

Instead, the *Hoxd11*-*Hoxd12* inversion also inverted a CTCF site, which had remained untouched in the two single gene inversion alleles. This site seems to be of particular importance as it is the first of a series of CTCF sites located at the 5’ extremity of the gene cluster that all display the same orientation i.e. opposite to those sites found at more central positions within the *HoxD* locus. This very region where the polarity of CTCF sites is inverted coincides with the strongest boundary effect that separates T-DOM from C-DOM, i.e. the two TADs flanking the gene cluster, likely due to large loops established in either direction (Rodriguez-Carballo et al. 2017). As a consequence of the *Hoxd11-Hoxd12* inversion, this site was relocated towards a more 3’ position with an opposite orientation. Therefore, rather than separating *Hoxd13* from the rest of the gene cluster, the new distribution of CTCF sites predicted a shift of the boundary upstream *Hoxd13* where two CTCF sites still have the requested orientations to induce a boundary effect.

Several observations support this interpretation. First, the *Hoxd13lacZ* reporter gene, which before the inversion mostly contacted a H3K27me3-labelled, negative domain on the centromeric side of the TAD boundary (including *Evx2*), contacted the telomeric side of the boundary after the inversion, in particular the inverted *Hoxd11* gene. Also, in the inverted configuration, *Evx2* remained robustly covered by H3K27me3 marks, suggesting that the new TAD boundary, at least in metanephros, had been moved between *Evx2* and *Hoxd13*. Finally, the inverted CTCF site continued to impose a boundary effect (as the CTCF site starting the series with the same orientation), yet this effect was local and did not prevent *Hoxd13lacZ* to contact the telomeric side of the *HoxD* cluster where potential metanephros enhancers were assumed to be.

### Deletion of the CTCF site

Because of the apparent importance of this particular CTCF binding site in preventing *Hoxd13* to respond to metanephros enhancers, we deleted it from a wild-type genetic background. This deletion lead to an up-regulation of *Hoxd13* in developing metanephros, supporting the proposal that the series of CTCF sites with a telomeric orientation is necessary to block potentially deleterious contacts between *Hoxd13* on the one hand, and ‘anterior’ enhancers on the other hand. In fact, this specific CTCF site maps exactly at the position where an elusive ‘polar silencer’ had been previously positioned based on a genetic approach (Kmita et al. 2000). The gain-of-expression was nevertheless transitory, likely due to the presence of the other CTCF sites that make this boundary very resilient (Rodriguez-C.) and *Hoxd13* transcripts were rapidly down regulated, which might account for the apparent absence of renal phenotype in these animals.

However, a slightly more pronounced and stable presence of *Hoxd13* mRNAs in the developing metanephros, as induced by the de-sequestering of the *Hoxd13* locus following a large centromeric inversion, lead to severe alterations in metanephros morphology already in heterozygous specimen. While such a correlation between the ectopic presence of *Hoxd13* transcripts and metanephros malformation or even agenesis was previously observed, the causative involvement of these transcripts in triggering the abnormal phenotype was never shown. Here we demonstrate that a mutation in the DNA binding region of the HOXD13 protein in *cis* with the (*Nsi-Itga6*) inversion is sufficient to entirely rescue the phenotype. This result indicates that HOXD13 is indeed responsible for the observed alterations and that this dominant negative effect is likely mediated by DNA binding.

### The logic of global *Hox* gene regulation

These latter results demonstrate the critical importance of insulating this gene from telomeric regulatory influences and thus help understand the general logic of the global regulation occurring at *Hox* loci. *Hox13* genes are generally positioned at the last position within vertebrate *Hox* clusters and, as such, they are the last to be activated and at the most rostral positions. Due to the dominant negative effect of their proteins over other *Hox* functions (posterior prevalence, see (Duboule and Morata 1994), their activation participates to the termination of various axial systems such as the limbs, the intestinal tract and the major body axis, as seen by various gain- and loss-of-function approaches (Economides et al. 2003; Young et al. 2009; Aires et al. 2019). Therefore, while it is essential that these genes remain associated with their respective *Hox* clusters such that there are activated in a coordinated spatial and temporal manners, they must be prevented to be transcribed too early and too anteriorly, which would have dramatic effects for the developing body. We believe this is the reason behind the evolution of such strong and resilient boundaries within *Hox* clusters, as illustrated by the unusually high concentration of CTCF sites.

Along with the emergence of global enhancers located on either side on the gene cluster, progressively forming these complex regulatory landscapes, these CTCF sites were used to allocate various sub-groups of *Hox* genes to particular telomeric (anterior, early) enhancers (Narendra et al. 2016), yet always by respecting the insulation of *Hox13* genes. This was shown for proximal limb enhancers (Andrey et al. 2013), for the intestinal caecum (Delpretti et al. 2013) and for the developing metanephros in this work. In this view, the bimodal gene regulatory system proposed by Andrey et al. (Andrey et al. 2013) and based on the presence of two TADs flanking the *HoxD* cluster illustrates the general logic of *Hox* gene regulation. On the one hand a positive ‘morphogenetic potential’ for the primary body axis, i.e. the ancestral structure where *Hox* genes’ functions were initially deployed, to organize the body plan by regulating the ‘rostral’ side of the *Hox* gene clusters. On the other hand, a negative inhibitory function for those regulations located in the other side and acting upon the termination of this genetic system by activating *Hox13* genes. Such a bi-modal regulatory strategy subsequently constrained and guided the evolution of regulatory innovations on either side of the cluster, along with the emergence of vertebrate morphological novelties.

## Materials and Methods

### Mouse stocks

The *HoxD*^*inv(11)*^, *HoxD*^*inv(12)*^, *HoxD*^*inv(11-12)del(11)d13lacZ*^ and *HoxD*^*del(CTCF:d12d13)*^ were obtained through the CRISPR/Cas9 technology using mouse zygotes. gRNAs were designed either to flank the locus of interest or to directly target a CTCF binding site. The gRNAs are listed in Supplementary Table 1. Each guide was cloned into the pX330:hSpCas9 (Addgene ID 42230) vector (Cong et al. 2013). The efficiency and specificity of these guides were evaluated *in silico* using either Cctop (Stemmer et al. 2015), CHOPCHOP (Montague et al. 2014) or crispr.mit.edu. The *HoxD*^*inv(11)*^ and *HoxD*^*inv(12)*^ alleles were obtained by pro-nuclear injection (PNI) of an equimolar solution of the two pX330:hSp:Cas9 vectors containing the appropriate guides (Mashiko et al. 2013). The *HoxD*^*inv(Nsi-Itga6)del(13HD)*^ allele was obtained in a similar fashion, except that the zygotes injected were obtained from a cross between *HoxD*^*inv(Nsi-Itga6)/+*^ male and wild-type female mice. The *HoxD*^*inv(11-12)del(11)d13lacZ*^ allele was obtained by electroporation in zygotes of a solution containing *Cas9* mRNA and the two desired gRNAs *in vitro* transcribed (Hashimoto and Takemoto 2015). The zygotes were obtained from a cross between *HoxD*^*inv(11-12)d13lacZ/+*^ males and wild-type females. The *HoxD*^*del(CTCF:d12d13)*^ allele was generated by electroporation of a solution containing a single gRNA *in vitro* transcribed, with *Cas9* mRNA within wild-type zygotes. The *d13lacZ*, *inv(11-12)d13lacZ* and *del(8-13)* alleles were described in Kmita et al., 2001 and Tarchini et al., 2005. The allele *inv(Nsi-Itga6)* was originally described in (Tschopp and Duboule 2011).

F0 animals were PCR genotyped at weaning and the breakpoints Sanger sequenced. These sequences were also used to reconstruct the different genomes used for alignments in this work. Positive F0 animals were back-crossed over wild-type (B6XCBA)F1 mice for at least two generations and single heterozygous F2 mutant animals were then used to expand each alleles. Transgenic integration of the pX330:hSp:Cas9 vectors was regularly observed in animals derived from pro-nuclear injection across different founders and alleles. The segregation of these transgenes was monitored by PCR and only pX330 negative animals were used to expand the new alleles. For PCR genotyping, tissue biopsies were lysed overnight at 56°C in lysis buffer (10mM KCl, 20mM TrisHCl pH 8.0, 10mM (NH4)2SO4, 1mM EDTA, 0.1% Triton,) complemented with 0.5% proteinase K (20mg/ml). After proteinase K heat-inactivation at 98°C for 10 minutes, PCR were performed with a standardized cycling protocol. The list of primers used for genotyping is provided in Supplementary Table 2.

### WISH and X-gal staining

WISH and X-gal staining were carried out using standard protocols. WISH were performed as in (Woltering et al. 2009) using DIG labelled RNA probes (*lacZ, Hoxd13*, *d12*, *d11*, *d10* and *lacZ*) described in (Dolle et al. 1991; Izpisua-Belmonte et al. 1991; Gerard et al. 1996; Zakany et al. 2001). For WISH performed on E13.5 metanephros, the abdominal cavity was opened and the embryo eviscerated to expose the metanephros before the ISH procedure. The duration of the proteinase K treatment was reduced to 6 minutes. *LacZ* detection by X-gal staining in metanephros at E13.5 was done on eviscerated embryos, fixed in 4% PFA for 20 minutes at 4°C. Staining was performed overnight at 37°C following standard X-gal staining procedure except that the staining solution was complemented with TrisHCl pH=7.4 (20uM final concentration) to avoid background signal in the metanephros. Pictures of specimen were taken with a DP74 camera coupled to an MVX10 binocular loop (Olympus). WISH pictures of *HoxD*^*inv(11)*^ and *HoxD*^*del(CTCF:d12d13)*^ embryos were reconstructed on their Z-axis using the software Helicon Focus (7.0.2) to facilitate the visualization of *Hoxd* genes expression patterns across different area of the embryo.

### Water consumption

Water intake was quantified for each genotype investigated by daily measurement of the weight of water bottles during a series of 6 consecutive days. At least three cages housing each single adult female mouse were used for each allele. Individual water consumption was then normalized by the corresponding average body mass recorded 3 times during the experiment.

### Histology

Metanephros were dissected from heterozygous adult mice, rinsed abundantly in 1x PBS and fixed overnight in 4% PFA prior to dehydration and paraffin embedding. Sections were then produced and stained at the HCF platform (EPFL).

### Genomic data

The NGS datasets were all mapped on GRCm38/mm10. The various mutated chromosomes 2 present in the *HoxD*^*inv(11)*^, *HoxD*^*inv(12)*^, *HoxD*^*d13lacZ*^, *HoxD*^*inv(11-12)d13lacZ*^ and *HoxD*^*del(CTCF:d12d13)*^ alleles were each reconstructed from Sanger sequencing of all the different breakpoints, including the *lacZ* reporter when relevant. The GTF annotations used in this work derive from ENSEMBL GRCm38.90 and are filtered against read-through/overlapping transcripts, keeping only transcripts annotated as ‘protein-coding’ for ‘protein-coding’ genes, thus discarding transcripts flagged as ‘retained_intron’, ‘nonsense-mediated decay’ etc., to conserve only non-ambiguous exons and avoid quantitative bias during data analysis by STAR/Cufflinks (Amândio et al. 2016). The mutant chr2 sequences and their associated GTF files are available on figshare (Darbellay et al. 2019). ChIP-seq and RNA-seq libraries were build and sequenced by the UniGe Genomics platform. ChIPmentation libraries were sequenced at GECF (EPFL). Bioinformatic analysis were done using our Galaxy server (the Bioteam Appliance Galaxy Edition, https://bioteam.net/products/galaxy-appliance, (Afgan et al. 2016). The 4C-seq datasets were analyzed with the HTSstation interface (BBCF, EPFL, (David et al. 2014)). Figures presented in this work were built using a genome browser assembled as an R script.

### RNA-seq

RNA was extracted from micro-dissected tissues using RNeasy plus micro kit (Qiagen) following the manufacturer recommendations. At least 1.5ug of total RNA (RIN>9) was prepared for stranded RNA sequencing using TruSeq kits (Illumina). Samples were poly(A)-enriched. Following sequencing, Illumina TruSeq adaptors were cleaved from the raw fastq files using Cutadapt (1.6) with the following settings -a AGATCGGAAGAGCACACGTCTGAACTCCAGTCAC (Martin 2011). Reads were then mapped to their respective genome using STAR (2.5.2b) with the ENCODE settings (Dobin et al. 2013). FPKM values were determined by Cufflinks (2.2.1) with the following options: -I 600000 -F 0.05 -j 0.05—compatible-hits-norm–multi-read-correct-library-type fr-firststrand -m 45 -s 20 --min-intron-length 40 (Trapnell et al. 2010). Normalized FPKM were then computed by determining coefficients extrapolated from a set of 1000 house-keeping stably expressed genes, in between the series of compared RNA-seq datasets. These coefficients were then applied to the respective FPKM values. Strand specific coverage was computed from uniquely-mapped reads and normalized to the number of million uniquely mapped reads for each RNA-seq dataset. Biological duplicates were produced for the *HoxD*^*del(CTCF:d12d13)*^ and *HoxD*^*inv(Nsi-Itga6)*^ datasets.

### RT-qPCR

RNAs were extracted from metanephros dissected from individual E18.5 fetuses and adult mice of equivalent age using RNeasy plus micro kit (Qiagen) following the manufacturer recommendations. For each sample, 1ug of total RNA was reverse transcribed using SuperScript VILO cDNA Synthesis Kit (ThermoFisher). RT-qPCR was performed in technical triplicates on a CFX96 real-time system (BIORAD) using the GoTaq qPCR Master Mix (Promega). Each RT-qPCR was carried out with at least two biological replicates and *ΔCt* were computed using the house-keeping gene *Rps9*. Primers’ sequences for RT-qPCR are listed in Supplementary Table 3.

### ChIP-seq and ChIPmentation

ChIP and ChIPmentation datasets were produced using the following commercial antibodies: 4ul anti-H3K4me3 (17-614, Millipore), 4ul anti-H3K27me3 (17-622, Millipore), 5ug anti-CTCF (61311, Active Motif) and 5ug anti-RAD21 (ab992, Abcam). ChIP were produced from 5 pairs of metanephros micro-dissected from E13.5 embryos (approx. 1x10^6^ cells). Pairs of metanephros were individually fixed in 1% Formaldehyde for 20 minutes at room temperature and stored at -80°C. Genotyping was done on tail biopsies. Shearing was performed on a Bioruptor (Diagenod) in 1% SDS. ChIP was done by over-night incubation of protA/protG magnetic beads conjugated with the desired antibody in 1.2ml dilution buffer, rotating at 30 rpm 4°C. Washes were in 1ml of washing buffers, incubated at 4°C for 2 minutes: 2x RIPA/2x RIPA-500mM NaCl/2x LiCl/2x TE. Beads were eluted and chromatin fixation was reversed during an over-night incubation at 65°C in presence of proteinase K. The eluate was treated with RNase A, PCI purified and precipitated. DNA was then quantified for TruSeq library construction.

The ChIPmentation protocol was adapted from (Schmidl et al. 2015). ChIPmentation datasets were produced from 4 pairs of metanephros (approx. 7.5x10^5^ cells) or from a fraction of forebrain micro-dissected from E13.5 embryos. Metanephros were dissociated for 10 minutes at 37°C, 700 rpm, in 400ul of a solution containing 1x PBS complemented with 10% FCS and 5ul Collagenase 10mg/ml (sterile filetered, C1764 Sigma). Fixation was done in a final concentration of 1% FA (w/o methanol, 28908 Thermo) for 10 minutes at room temperature and cells were then stored at -80°C. Genotyping was done on brain or tail biopsies. Chromatin shearing was done using a Covaris E220 in 120 ul sonication buffer (0.25% SDS) with the following settings: duty cycle at 2%, peak incident power at 105W, water level set at 6 with the amplifier engaged and a treatment of 8 minutes. The volume for the over-night IP (4°C, 25 rpm) was adjusted at 300 ul using a dilution buffer to lower the SDS concentration and buffer the pH (final concentration of 0.1% SDS, 20 mM HEPES). Following two LiCl wash, magnetic beads were then washed 2x in 1ml 10uM TrisHCl pH=7.4 and finally resuspended in 24 ul of pre-warmed tagmentation buffer (FC-121-1030, Illumina). 1 ul of TDE1 enzyme was added to the mix and the beads were incubated for 120” at 37°C. DNA purification, quantification and Nextera library construction were done following the published ChIPmentation protocol (Schmidl et al., 2015) using indexed Illumina adapters (FC-121-1011, Illumina). Illumina adaptors were removed from the ChIP and ChIPmentation reads using Cutadapt (1.6) with the following settings; for TruSeq (single-end reads) -a AGATCGGAAGAGCACACGTCTGAACTCCAGTCAC and for Nextera (pair-end reads) -a CTGTCTCTTATACACATCTCCGAGCCCACGAGAC for R1 and -a CTGTCTCTTATACACATCTGACGCTGCCGACGA for R2 (Martin 2011). The single-end or pair-end reads were then mapped to the corresponding genome using Bowtie2 (2.2.6.2) with default parameters (Langmead and Salzberg 2012). Reads were filtered for a MAPQ≥30 and the final coverage was obtained after duplicates removal, estimation of the average fragments size and extension of the reads by MACS2 (2.1.0.20151222, (Zhang et al. 2008).

### 4C-seq

4C-seq datasets were generated using pairs of metanephros individually micro-dissected from E13.5 embryos. Samples were then dissociated for 10 minutes at 37°C, 700 rpm, in 250ul of a solution containing 1x PBS complemented with 10% FCS and 5ul Collagenase 10mg/ml (sterile filtered, C1764 Sigma). Fixation was done in a final concentration of 2% PFA for 10 minutes at room temperature. Following glycine quenching and two 1x PBS washes to stop the fixation, the cells were then lysed and nuclei stored at -80°C. Genotyping of the corresponding samples was done on brain or tail biopsies processed in parallel. Pools of 20-25 pairs of metanephros of similar genotypes were then used to produce a double-digest libraries following the protocol described in (Noordermeer et al. 2011), using *NlaIII* and *DpnII*. For each 4C-seq viewpoint, 11 individual PCR were run and pooled together, using between 100-150 ng of the double-digested initial library per single PCR. Series of viewpoints were then multiplexed together and sequenced in a HiSeq2500. The reads were demultiplexed, mapped over mutant genomes, normalized and analyzed using the workflow provided by HTSstation (David et al. 2014). The profiles were smoothened using a window size of 11 fragments, and the final coverage was normalized considering the total signal spanned 5Mb on the two sides of each viewpoint. 4C-seq primers used in this work are listed in Supplementary Table 4.

### Animal experimentation

All experiments were performed in agreement with the Swiss law on animal protection (LPA), under license No GE 81/14 (to DD).

### Accession Numbers

RNA-seq, 4C-seq, ChIP-seq and ChIPmentation datasets are available from the NCBI Gene Expression Omnibus repository under the reference GSE127870.

## Acknowledgements

We thank Dr. A. Necsulea for help with bioinformatic analyses; S. Gitto and T.-H. Nguyen Huynh for technical help; the Geneva Genomics Platform (University of Geneva), the Gene Expression Core Facility and the Transgenic Core Facility at CPG as well as the Histology Core Facility (Ecole Polytechnique Fédérale in Lausanne). We are grateful to Dr. L. Beccari for sharing with us a pair of 4C-seq primers (*ilacZ*). We also thank present and past members of the Duboule laboratories for discussions, sharing reagents and protocols. This work was supported by funds from the Ecole Polytechnique Fédérale in Lausanne, the University of Geneva, the Swiss National Research Fund, European Research Council grants Regul*Hox*, and the Claraz Foundation (to D.D).

## Author Contributions

Conceptualization: Fabrice Darbellay, Denis Duboule.

Formal analysis: Fabrice Darbellay, Lucille Lopez-Delisle.

Funding acquisition: Denis Duboule.

Investigation: Fabrice Darbellay, Célia Bochaton, Jozsef Zakany, Bénédicte Mascrez, Patrick Tschopp, Saskia Delpretti

Methodology: Fabrice Darbellay, Lucille Lopez-Delisle, Jozsef Zakany, Bénédicte Mascrez

Project administration: Denis Duboule.

Resources: Denis Duboule.

Supervision: Denis Duboule.

Validation: Denis Duboule.

Writing – original draft: Fabrice Darbellay, Denis Duboule.

Writing – review & editing: All authors.

---

**Supplementary Figure 1:**
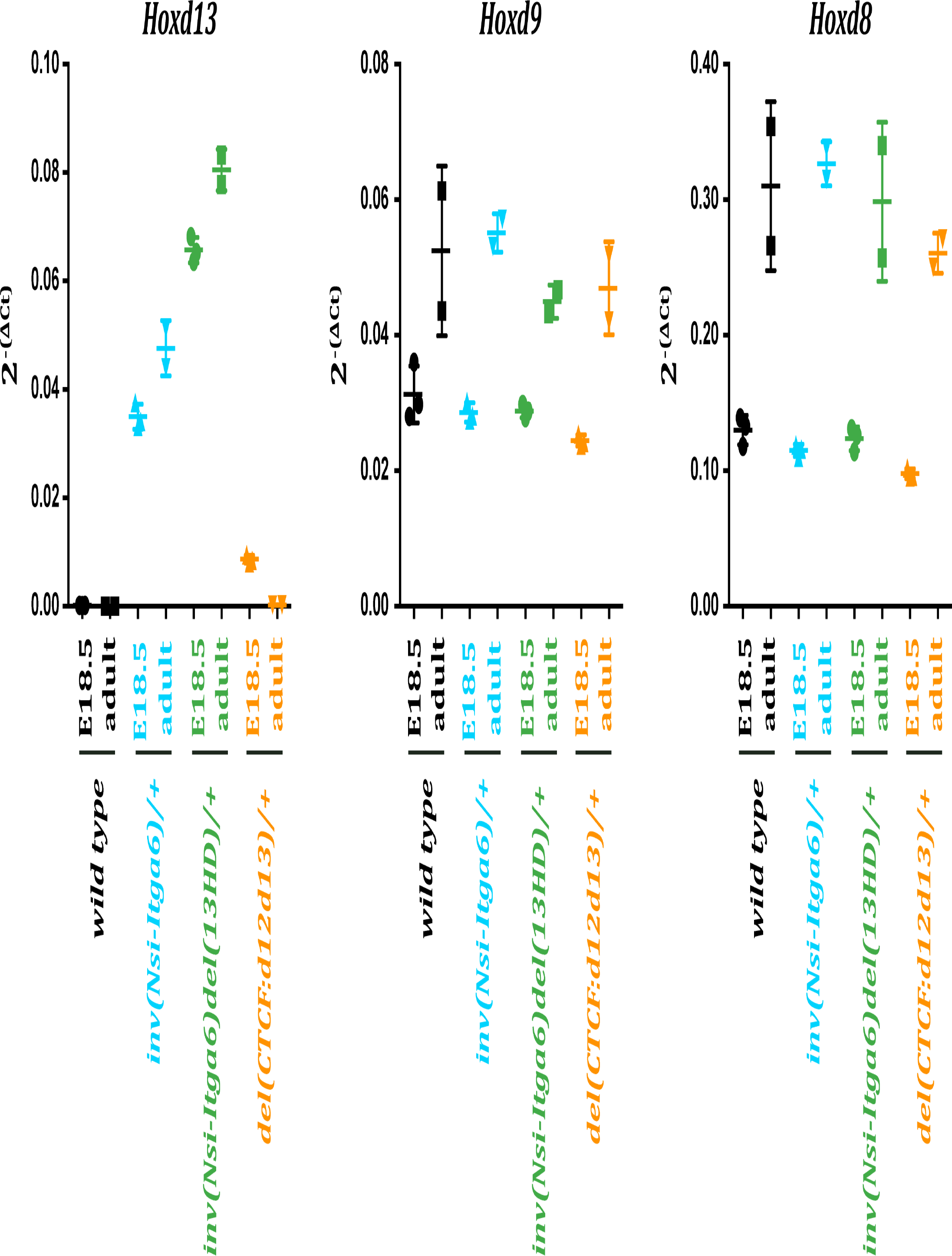
RT-qPCR performed on RNAs extracted from E18.5 and adult mice metanephros revealed the transient nature of the gain-of-expression of *Hoxd13* in metanephros of the *del(CTCF:d12d13)* mutant (orange) as compared to the *inv(Nsi-Itga6)* (blue) and *inv(Nsi-Itga6)del(13HD)* (green) alleles. No signal was recorded in the wild-type control (black), in agreement with the absence of any significant *Hoxd13* transcription at these two different time points. *Hoxd9* and *Hoxd8*, two genes which normally remain transcribed in late embryonic and adult metanephros do not display any significant change in their relative expression levels across these four genotypes. *Rps9* was used as an internal normalizer in these series to compute *ΔCt*. Mutant metanephros were dissected from heterozygous animals. Three (E18.5) and two (adult) biological replicates were used to generate this series.

**Supplementary Figure 2:**
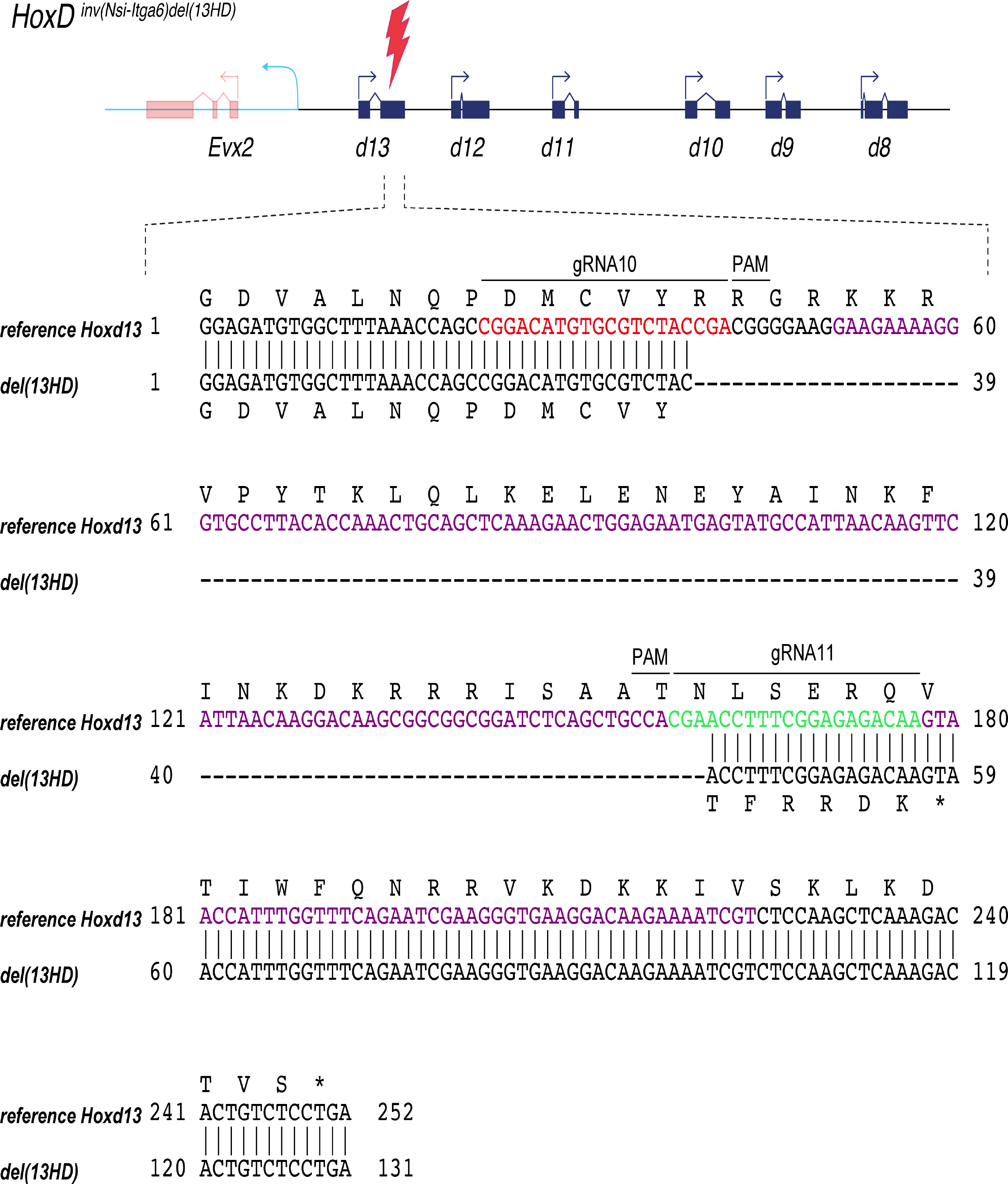
The loss-of-function of *Hoxd13* in the *HoxD*^*inv(Nsi-Itag6)del(13HD)*^ mutant (red lighting) was produced by a CRISPR/Cas9 mediated deletion of 120 bp in the homeobox of *Hoxd13* (purple), positioned in the second exon of this gene. This deletion was obtained by the combined action of the guides gRNA10 (red) and gRNA11 (green), in *cis* with the *inv(Nsi-Itga6)*. The *del(13HD)* mutation results in a truncated HOXD13 protein, lacking a functional DNA-binding homeodomain.

**Supplementary Table 1:**
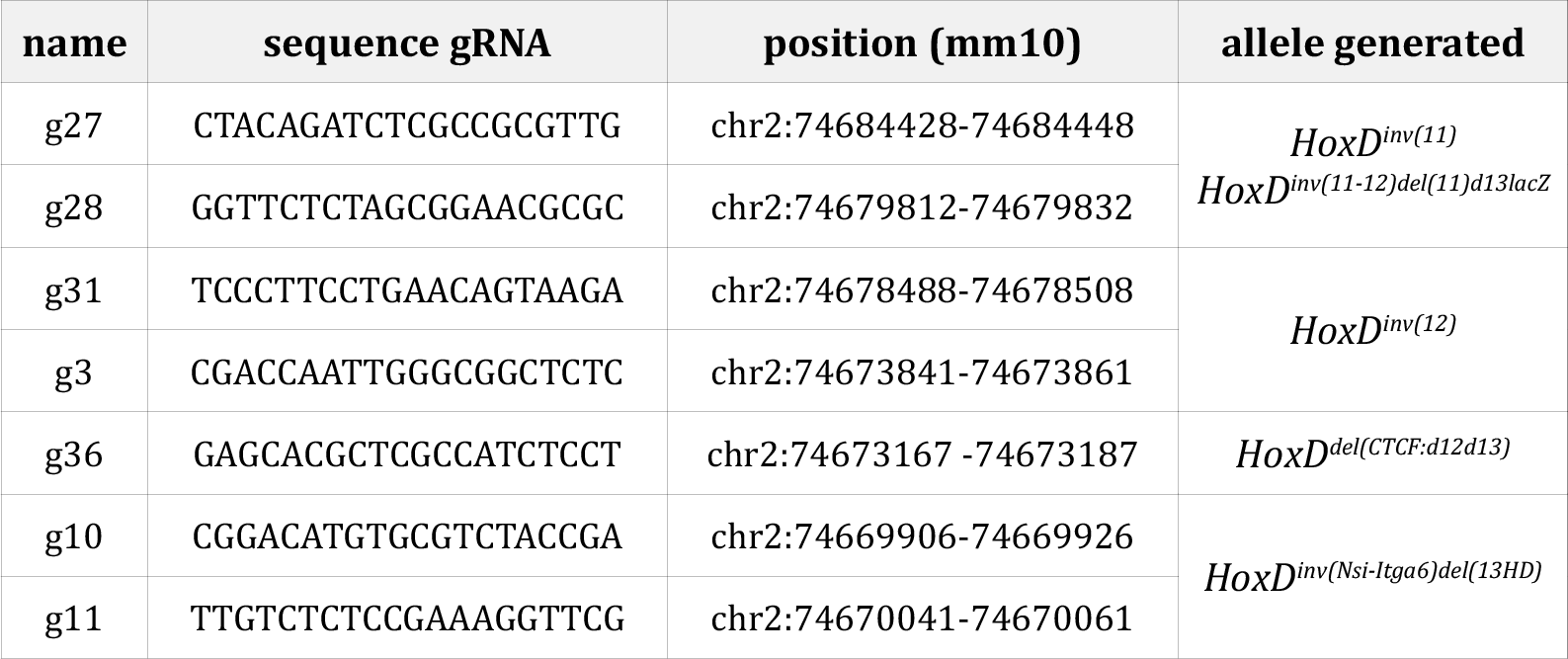
Sequence and position of gRNAs used in this work

**Supplementary Table 2:**
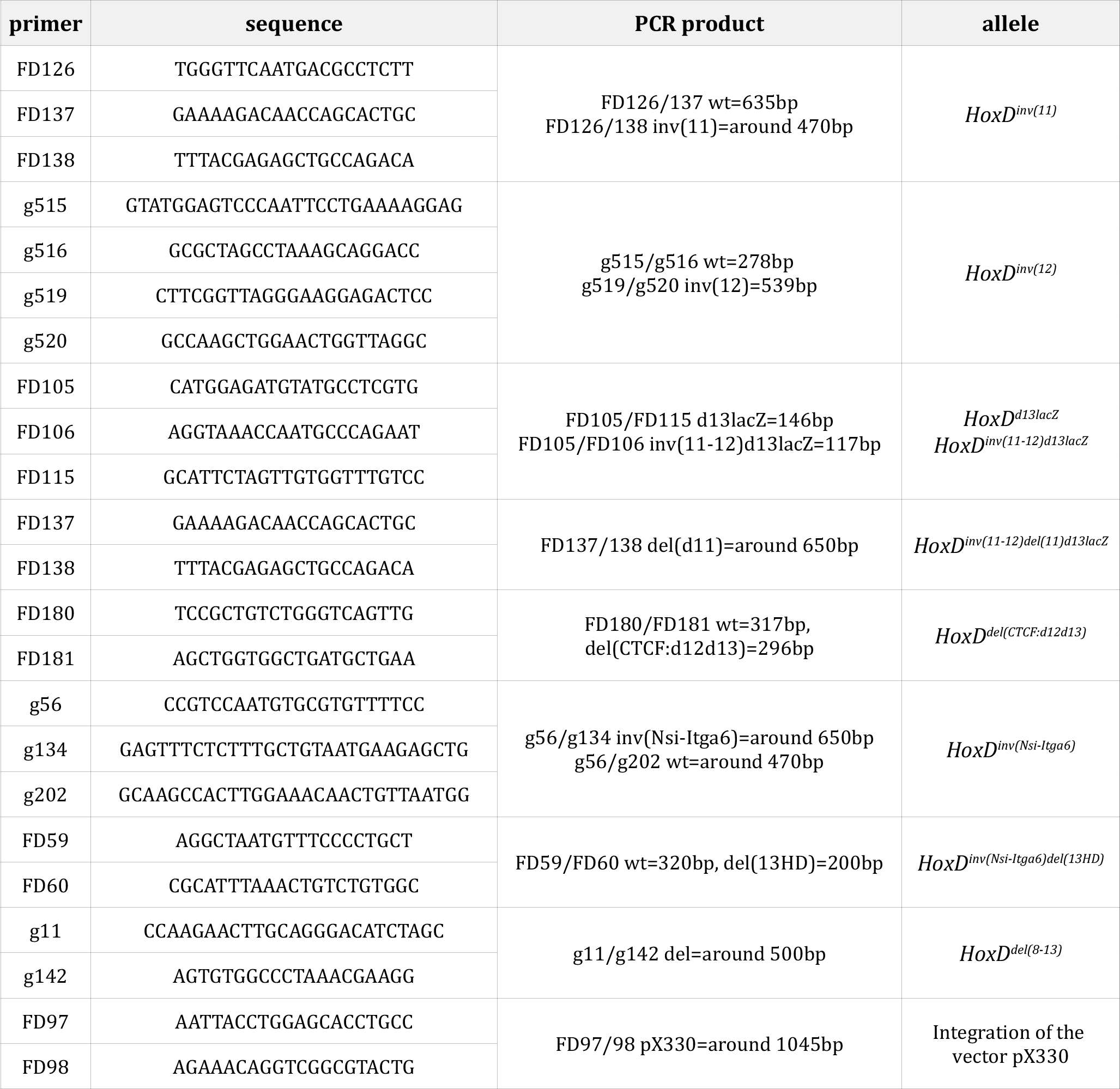
Sequence of genotyping primers used in this work

**Supplementary Table 3:**
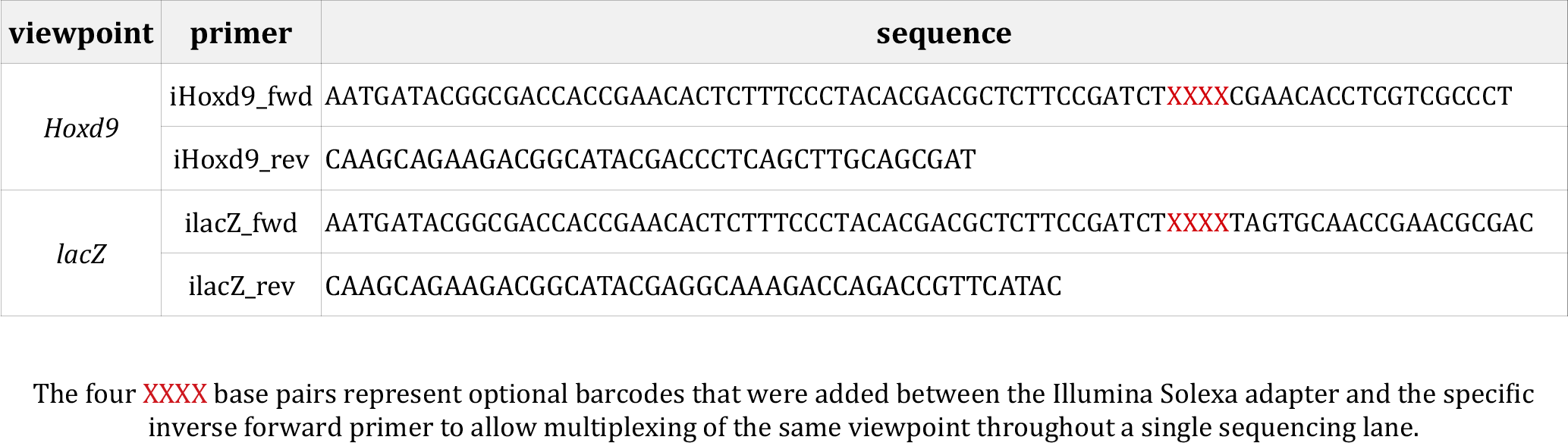
Sequence of 4C-seq primers used in this work

**Supplementary Table 4:**
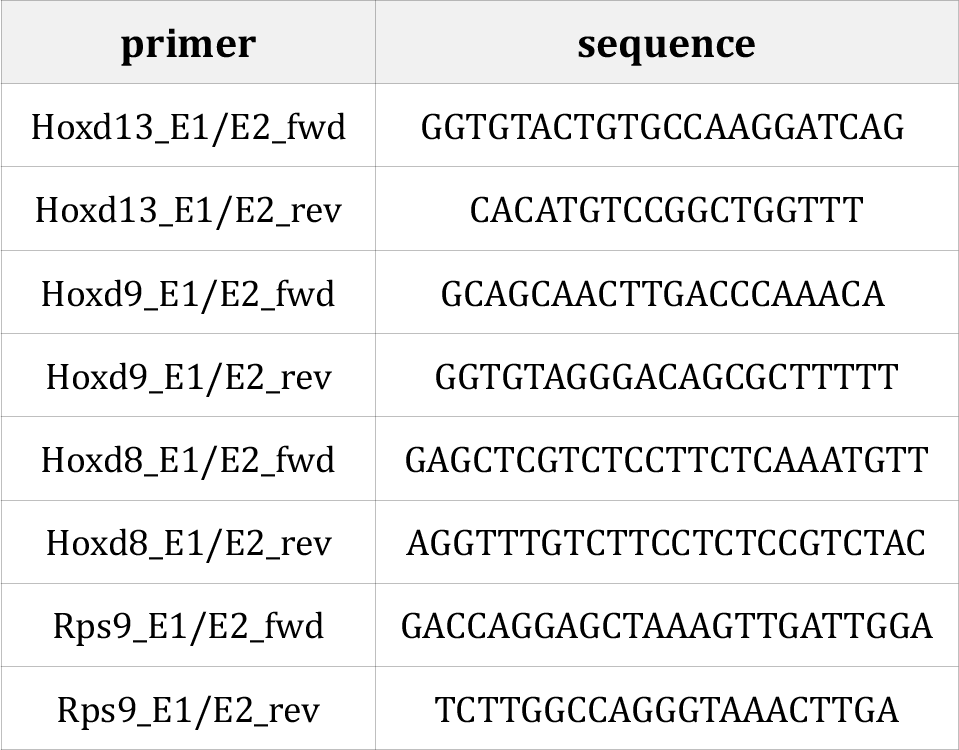
Sequence of RT-qPCR primers used in this work

